# Single cell transcriptomics unveiled that early life BDE-99 exposure reprogrammed the gut-liver axis to promote a pro-inflammatory metabolic signature in male mice at late adulthood

**DOI:** 10.1101/2023.06.22.546158

**Authors:** Joe Jongpyo Lim, Michael Goedkin, Yan Jin, Haiwei Gu, Julia Yue Cui

**Author notes:** Address for correspondence: Julia Yue Cui, PhD, DABT Associate Professor Department of Environmental and Occupational Health Sciences University of Washington 4225 Roosevelt Way NE, Seattle, WA 98105.

## Abstract

Polybrominated diphenyl ethers (PBDEs) are a class of legacy flame retardants that bioaccumulate in the environment, raising global health concerns. The gut microbiome is an important regulator of liver including xenobiotic biotransformation, nutrient homeostasis, and immune regulation. Using bulk RNA-Seq, we recently showed that neonatal exposure to BDE-99, a human breast milk-enriched PBDE congener, up-regulated pro-inflammation- and down-regulated drug metabolism-related genes predominantly in males in young adulthood. However, it remains unknown whether such dysregulation persists into late adulthood, how various cell types in the liver contribute to the hepatotoxicity, and to what extent gut microbiome is involved in BDE-99 mediated developmental reprogramming of the liver. To address these knowledge gaps, male C57BL/6 mouse pups were orally exposed to corn oil (10 ml/kg) or BDE-99 (57 mg/kg) once daily from postnatal days 2-4. At 15 months of age, single cell transcriptomics (scRNA-seq) in liver showed that neonatal BDE-99 exposure down-regulated key xenobiotic- and fatty acid metabolizing enzymes and up-regulated genes involved in microbial influx in hepatocytes. Neonatal BDE-99 exposure also led to a persistent increase in the hepatic proportion of neutrophils, a predicted increase of macrophage migration inhibitory factor (MIF) signaling, which activates macrophage populations, as well as histopathological abnormalities of the liver in 15 months of age. The BDE-99 mediated hepatic reprogramming is associated with decreased intestinal tight junction protein (Tjp) transcripts, persistent dysbiosis of the gut microbiome, and dysregulation of inflammation-related fatty acid metabolites. ScRNA-seq in germ-free (GF) mice demonstrated the necessity of a normal gut microbiome in maintaining hepatic immunotolerance. Fecal microbiome transplant to GF mice using large intestinal microbiome from adults that were neonatally exposed to BDE-99 down-regulated Tjp transcripts and up-regulated several cytokines in the large intestine. In conclusion, neonatal BDE-99 exposure reprogrammed the cell type-specific gene expression and cell-cell communication networks in liver towards a pro-inflammation with compromised metabolic functions at late adulthood. Importantly, gut microbiome is necessary in promoting immunotolerance in the liver, and BDE-99-mediated pro-inflammatory signaling may be partly due to the dysregulated gut environment.

## INTRODUCTION

The developmental origins of health and disease (DOHaD) hypothesis emphasizes the profound influence of early life exposures on health and diseases later in life (Mandy and Nyirenda, 2018). There is a sensitive developmental time window for exposures to toxic environmental chemicals that may have a life-long impact on risks of complex diseases, such as obesity, type II diabetes, and tumorigenesis (Stein *et al*., 2019; Van den Bergh, 2011; Filbin and Monje, 2019; Lacagnina, 2020). For example, early life exposure to endocrine-disrupting chemicals, such as bisphenol A (BPA) increased the incidence of liver tumors in adult mice (Weinhouse *et al*., 2014). In addition, a recent study showed that prenatal exposure to Polybrominated diphenyl ethers (PBDEs) is associated with liver injury in children (Midya *et al*., 2022). However, very little is known regarding the underlying mechanisms. Polybrominated diphenyl ethers (PBDEs) are legacy flame-retardants that were used in a wide variety of consumer products, such as electrical equipment, construction materials, coatings, textiles, and polyurethane foam (Siddiqi *et al*., 2003). As persistent organic pollutants, PBDEs bio-accumulate in the environment and are detected in household dust, fish, poultry, water, and human breast milk (Bocio *et al*., 2003; Xu *et al*., 2019; Imm *et al*., 2009; Gascon *et al*., 2012). In both animal models and humans, acute and chronic exposure to PBDEs are linked to a wide range of diseases, such as thyroid hormone disorders, neurotoxicity, hepatic oxidative stress, and carcinogenesis (Allen *et al*., 2016; Dorman *et al*., 2018; Manuguerra *et al*., 2019; Dunnick *et al*., 2018). Despite the production ban in the United States in 2004, the concern about exposure to PBDEs continues due to their persistent and bio-accumulative nature (Varshavsky *et al*., 2020).

The liver is a critical organ for xenobiotic biotransformation and nutrient homeostasis (Trefts *et al*., 2017). The global incidence of liver diseases, such as type 2 diabetes-associated nonalcoholic fatty liver diseases, liver fibrosis, and liver cancer, has been steadily increasing (Cheemerla and Balakrishnan, 2021; Asrani *et al*., 2019; Tolman *et al*., 2007). The liver contains various cell types with distinct roles (Fig. 1A). Hepatocytes are important for drug metabolism and transport as well as intermediary metabolism. Cholangiocytes provide the structure of bile duct. Kupffer cells, which are the resident macrophages in the liver, regulate the innate immune system. Endothelial cells line the hepatic sinusoids. Stellate cells store vitamin A, and myofibroblasts (i.e. activated stellate cells) participate in regulating the hepatic immunological processes. Other immune cell types, such as B and T cell populations, monocytes and monocyte-derived macrophages (MDMs), dendritic cells, and neutrophils may be recruited to the liver (Crispe, 2003; Zhao *et al*., 2020; Wen *et al*., 2021) to facilitate immune response. Environmental-stressor-mediated dysregulation of the proportions of various hepatic cell types and cell-cell communications may predispose the onset of various liver diseases.

**Figure 1.**
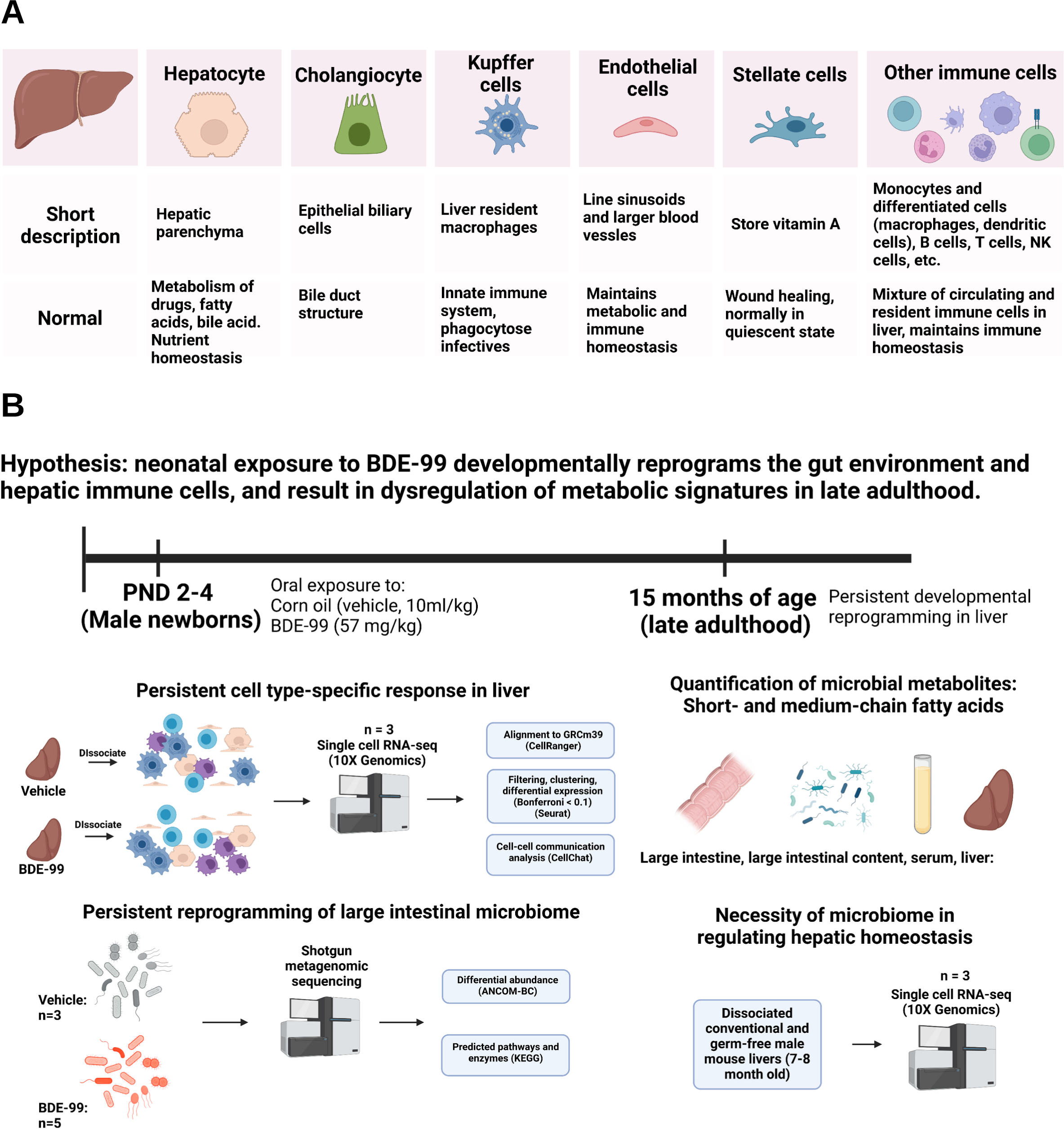
**A.** Summary of known cell types detected in liver and their functions during normal and diseased states. **B.** Experimental design, data analysis, and key findings. From PND 2-4, male C57BL/6 mouse pups were supralingually exposed to BDE-99 (57 mg/kg) or corn oil (10 ml/kg) as the vehicle control. Livers were removed at 15 months of age for scRNA-Seq (n=3 per group). The shift in expression in drug processing signatures, cell-cell communication patterns, and immunological markers were investigated in each cell type. For mechanistic investigations of the gut-liver axis in mediating PBDE hepatotoxicity, we compared the hepatic transcriptomic signatures to our recently published dataset on gut microbiome in adult male C57BL/6 mice that were developmentally exposed to BDE-99 or vehicle using the same dosing regimen. Furthermore, the necessity of the gut microbiome in maintaining basal hepatic immune tolerance is validated using scRNA-seq conducted in livers of adult conventional and germ-free mice.

Inflammation is an important contributor to various liver diseases (Del Campo *et al*., 2018; Tanwar *et al*., 2020). Chronic liver inflammation increases the risks of the development of fibrosis and cirrhosis, which are the 12^th^ leading cause of death in the United States (Koyama and Brenner, 2017). Various environmental stressors such as environmental toxicants, pathogenic microbes, microbial constituents, oxidative stress, as well as metabolic disorders such as insulin resistance and lipotoxicity can activate various pro-inflammatory cytokines to promote liver injury (Del Campo *et al*., 2018). Inflammation involves multiple cell types in the liver: chronic inflammation can activate hepatic stellate cells, which undergo trans-differentiation to become myofibroblasts – the main extra-cellular matrix-producing cells in the liver, and this subsequently contributes to liver fibrosis and cirrhosis (Tanwar *et al*., 2020). Kupffer cells are a significant source of chemoattractant molecules for T-cells and are also implicated in the pathogenesis of various liver diseases such as viral hepatitis, steatohepatitis, alcoholic liver disease, intrahepatic cholestasis, rejection of liver transplant, and liver fibrosis (Kolios *et al*., 2006). While Kupffer cells are the resident macrophages in the liver, other inflammatory cells such as neutrophils, dendritic cells (DCs), T-cells, and infiltrating macrophages also contribute to liver inflammation (Koyama and Brenner, 2017). In addition, Hepatocytes can produce pro-inflammatory mediators, including cytokines, chemokines, adhesion molecules, and other proteins that influence immune cell levels and function (Allen *et al*., 2011). An important mechanism of PBDE-mediated toxicity is inflammation. National Health and Nutrition Examination Survey (NHANES) data from 2003-2004 showed that serum PBDE concentrations in US human samples had a positive trend of association with liver injury and inflammation markers (Yuan *et al*., 2017). Therefore, it is important to understand the cell type-specific expression of inflammation-related genes as well as the cell-cell communications within the inflammasome complex to better understand the mechanisms of liver diseases.

One important component that regulates hepatic homeostasis is the gut environment. Via the portal vein, the liver receives the majority of its blood supply that passes through the gut (Kalra *et al*., 2022). Communication factors, such as gut microbial metabolites, microbial fragments, and other signaling molecules enter the hepatic environment, where they are processed and influence hepatic homeostasis (Ralli *et al*., 2022). Importantly, the disturbance of the intestinal barrier may indicate altered gut microbial composition, as well as their metabolic outputs, and an increased influx of these products to the liver (Genua *et al*., 2021; Stolfi *et al*., 2022). Recently, using bulk-RNA-Seq, we showed that early life exposure to BDE-99, a human breast milk-enriched PBDE congener, persistently reprograms the hepatic transcriptome at postnatal day (PND) 60 in mice, with males being more susceptible than females (Lim *et al*., 2021). Neonatal exposure to BDE-99 up-regulated hepatic immune response signatures and down-regulated genes involved in xenobiotic biotransformation in PND 60 males. This was associated with increases in distinct short-chain fatty acids (SCFAs) and related gut microbes that have the production capacity of the fatty acids. Together, these results suggest that early-life PBDE exposure altered the liver to a neoplastic-like state from epigenetic reprogramming of the liver, which is associated with a perturbed gut microbiome. However, it remains unknown whether the BDE-99 mediated developmental reprogramming of the gut-liver axis persists into late adulthood, how various hepatic cell types and their interactive networks are developmentally reprogrammed, and whether the perturbed gut microbiome mechanistically contributes to BDE-99 mediated hepatotoxicity. Therefore, in the present study, we tested our hypothesis that early life exposure to BDE-99 reprograms immune cells and hepatocytes in the developing liver to promote inflammation and reduce xenobiotic biotransformation capacities in late adulthood. We also tested to what extent the BDE-99-mediated dysbiosis of the gut microbiome is involved in the developmental reprogramming of the gut-liver axis.

## MATERIALS AND METHODS

**(note: the overall experimental design is summarized in Fig. 1B)**

### Chemicals and dosing regimen

2,2’,4,4’,5-pentabromodiphenyl ether (BDE-99 [CAS No. 60348-60-9]) was purchased from AccuStandard, Inc. (New Haven, Connecticut). Corn oil was purchased from Sigma-Aldrich (St. Louis, MO). BDE-99 was dissolved in corn oil and filtered using a 0.22 μm Millipore Express Plus Membrane filter (EMD Millipore, Temecula, California). All mice used in this study were housed according to the Association for Assessment and Accreditation of Laboratory Animal Care International guidelines. The study was approved by the University of Washington Institutional Animal Care and Use Committee. Eight-week-old specific pathogen-free C57BL/6J mice were purchased from the Jackson Laboratory (Bay Harbor, Maine), and were acclimated to the animal housing facility at the University of Washington for at least three breeding generations. Mice were housed in standard air-filtered cages using autoclaved bedding (autoclaved Enrich-N’Pure, Andersons, Maumee, OH). Mice had *ad libitum* access to non-acidified autoclaved water, as well as standard rodent chow (LabDiet No. 5021 for breeding pairs or to LabDiet No. 5010 for weaned pups) (LabDiet, St. Louis, MO). From postnatal day (PND) 2 to PND 4, male pups were supralingually exposed to BDE-99 (57 mg/kg, n = 5) or corn oil (vehicle control, 10 ml/kg, n = 3) once daily for three consecutive days. Litters and cages were randomly assigned to each exposure group. Pups were weaned at PND 21. At 15 months of age, serum, liver, small and large intestines, and fresh stool were collected. The remaining livers were subject to dissociation. To determine the role of the microbiome in regulating hepatic cell types, livers from 7∼8-month-old conventional (CV) or germ-free (GF) mice were collected.

### Whole liver dissociation

Fresh whole livers rinsed in Dulbecco’s phosphate-buffered saline (DPBS) (Cat# 14190136, Thermo Fisher Scientific, Waltham, MA) were minced to 1 – 3 mm pieces using surgical scissors. The minced livers were then placed in 15 mL conical tubes containing 10 mL of DPBS. Using serological pipettes, the DPBS was replaced by 10 mL of dissociation enzyme mixtures containing Liberase (Cat# 540115001, Sigma Aldrich, St. Louis, MO) and Dispase II (Cat# D4693-1G, Sigma Aldrich, St. Louis, MO) dissolved in Hanks’ Balanced Salt Solution (Cat# 14025-092, Thermo Fisher Scientific, Waltham, MA). The tubes with the livers and enzyme mix were incubated at 37 °C for 40 minutes using a Roto-Therm H2020 incubator (Benchmark Scientific Inc., Sayreville, NJ). After the incubation, the cells and un-dissociated fragments were strained using a 40-micron cell strainer. Strained cells were centrifuged at 300 g for 5 minutes at 4 °C and re-suspended in 5 mL red blood cell lysis buffer on ice for 8 minutes. Cells were centrifuged again at 300 g for 5 minutes at 4 °C. Dead and dying cells were filtered using the Dead Cell Removal Kit (Miltenyi Biotec, Cologne, Germany) following the manufacturer’s instructions. The viability was checked using a hemocytometer under a light microscope (Labophot-2, Nikon, Tokyo, Japan). Samples with greater than 85% viability were then cryopreserved using 10% Dimethylsulfoxide (Cat# BP231-100, Thermo Fisher Scientific, Waltham, MA) and 90% fetal bovine serum (Cat# F2442-50ML, Sigma Aldrich, St. Louis, MO) until further analysis.

### Single cell RNA sequencing

Cryopreserved cells were thawed using a water bath at 37 °C for 2 minutes, followed by serial dilution in DPBS until 32 mL was reached. Cells were centrifuged and re-suspended in DPBS until a concentration of approximately 100 cells/ μL was reached. The re-suspended cells (n=3), targeting 10,000 cells per sample, were then subject to scRNA-seq using a Chromium Next GEM single cell 3’ v3.1 kit and a Chromium X controller (10X Genomics, Pleasanton, CA) following the manufacturer’s instructions. The created libraries were then sequenced using the NovaSeq platform at paired-end 150 bp (∼11 M reads per sample).

### Data analysis of single cell RNA sequencing

Raw data were processed using the Cell Ranger v7.0 (10X Genomics, Pleasanton, CA). Processed data were read into R version 4.2.2 (R Core Team, 2022) for further analyses. Filtering and normalization were performed using the default parameters using Seurat v4 (Hao *et al*., 2021). Clustering was performed using the first 35 principal components, and the standard deviation obtained through principal component analysis was 1.5. Cell type labeling was performed using the differentially expressed genes using the function FindAllMarkers with default parameters in Seurat. Cell-cell communication analysis was done using the CellChat (v. 1.5) package (Jin, 2022). Gene ontology enrichment was done using the TopGO (v.2.48.0) package (Alexa A, 2022) using all detected genes as the background. Heatmaps were made using the ComplexHeatmaps (v. 2.13.1) package. All plots other than heatmaps were created using ggplot2 (v. 3.3.6).

### Liver Hematoxylin and Eosin staining

A fraction of the liver from each mouse was preserved in 4% formalin (Cat# SF100-4, Thermo Fisher Scientific, Waltham, MA) followed by 70% ethanol for histology. Hematoxylin and Eosin (H&E) staining was performed by the University of Washington Histology Core in the Department of Comparative Medicine. Briefly, all staining procedures were performed in a well-ventilated area with personal protective equipment. Liver slides were prepared, dried overnight, and placed in the oven at 60°C for approximately 30 minutes. The slides were stained in the following protocol: xylene for 5 minutes, rinsed in 100% ethanol for 4 minutes, 95% ethanol for 2 minutes, de-ionized (DI) water for 1 minute, hematoxylin for 3.5 minutes, DI water for 1 minute, clarifier for 30 seconds, DI water for 1 minute, bluing for 1 minute, DI water for 1 minute, eosin for 15 seconds, 95% ethanol for 1.5 minutes, 100% ethanol for 2 minutes, and xylene for 4 minutes. Images were then taken (Nanozoomer HT-9600, Hamamatsu Photonics, Japan), and histological incidence and severity scoring was performed by a board-certified veterinary pathologist.

### Metagenomic shotgun sequencing

DNA from large intestinal content was extracted using the EZNA Stool DNA kit (Omega Bio-Tek Inc., Norcross, GA). Shallow shotgun metagenomic sequencing was performed at 2 million reads (Diversigen, New Brighton, MN). DNA sequences were aligned to a curated database containing all representative genomes in RefSeq for bacteria with additional manually curated mouse-specific Metagenomically Assembled Genomes (MAGs) and cell-cultured genomes. Only high-quality MAGs (Completeness > 90% & Contamination < 5% via checkm) were considered. Alignments were made at 97% identity against all reference genomes. Every input sequence was compared to every reference sequence in the Diversigen DivDB-Mouse database using fully gapped alignment with BURST. Ties were broken by minimizing the overall number of unique Operational Taxonomic Units (OTUs). For taxonomy assignment, each input sequence was assigned the lowest common ancestor that was consistent across at least 80% of all reference sequences tied for best hit. Taxonomies are based on Genome Taxonomy Database (GTDB r95). Samples with fewer than 10,000 sequences were discarded. OTUs accounting for less than one-millionth of all strain-level markers and those with less than 0.01% of their unique genome regions covered (and < 0.1% of the whole genome) at the species level were discarded.

For functional analysis, Kyoto Encyclopedia of Genes and Genomes Orthology groups (KEGG KOs) were used with alignment at 97% identity against a gene database derived from the strain database used above (DivDB-Mouse). KOs were collapsed to level-2 (phylum) and −3 (class) KEGG pathways and KEGG Modules. Count tables for taxonomic, enzyme, and pathway data were transformed using the centered-log ratio (CLR) method (Quinn *et al*., 2019). Statistical testing for all count data was analyzed using ANCOM-BC2 (Lin and Peddada, 2020) with BH-FDR < 0.05. Significant taxa were plotted as heatmaps using the R package ComplexHeatmap (v.2.13.1). Bar plots were created using ggplot2 (v 3.3.6).

### Quantification of short-chain and medium-chain fatty acids

Short-chain and medium-chain fatty acids and their intermediate precursors were quantified as previously described (Gu *et al*., 2021; Dutta *et al*., 2022; Gomez *et al*., 2021). Briefly, approximately 50 mg of each tissue sample were homogenized in a mixture of 20 μl hexanoic acid-6,6,6-d_3_ (IS; 200 µM in H2O), 20 μl sodium hydroxide solution (NaOH, 0.5 M in water), and 480 μl methanol (MeOH). Then 400 μl of methanol was added. The pH of the mixture was adjusted to approximately 10. Samples were stored under −20°C for 20 min and then centrifuged at 21,694 × g for 10 minutes. A final volume of 800 μl of supernatant was collected. Samples were then evaporated to dryness, reconstituted in 40 μl of methoxyamine hydrochloride in pyridine (20 mg/ml), and stored at 60°C for 90 minutes. Subsequently, 60 μl of N-Methyl-N-tert-butyldimethylsilyltrifluoroacetamide was added and samples were heated to 60°C for 30 minutes. Samples were then vortexed for 30 seconds and centrifuged at 21,694 × g for 10 min. Finally, 70 μl of supernatant was collected from samples for gas chromatography-mass spectrometry (GC-MS) analysis.

GC-MS experiments were performed on an Agilent 7820A GC-5977B MSD system (Santa Clara, California) by injecting 1 µl of prepared samples. Helium was used as the carrier gas with a constant flow rate of 1.2 ml/min. The separation of metabolites was achieved using an Agilent HP-5ms capillary column (30 m × 250 × 0.25 µm). The column temperature was maintained at 60°C for 1 minute, and then increased at a rate of 10°C/minute to 325°C and held at this temperature for 10 min. The injector temperature was 250°C, and the operating temperatures for the transfer line, source, and quadruple were 290°C, 230°C, and 150°C, respectively. Mass spectral signals were recorded after a 4.9 min solvent delay. One-way analysis of variance (ANOVA) followed by Tukey’s post hoc test was performed for each metabolite in R for the analysis of liver metabolites (adjusted *p*-value < 0.05).

### Fecal microbiota transplantation

To investigate the functional role of changes in microbial composition in the large intestine, fecal microbiota transplant (FMT) was administered to 8-12 week-old male germ-free adult mice using the large intestinal content of adult conventional mice that were neonatally exposed to vehicle or BDE-99. The FMT procedure was based on previous publications (Turnbaugh *et al*., 2006; Ridaura *et al*., 2013). The intestinal content of donors was flushed out of the large intestine with sterile PBS. The intestinal content was then diluted to approximately 50 mg/mL in sterile PBS. Each sample was mixed with a 1 mL pipette thoroughly. Two hundred μl of the diluted homogenate was orally gavaged to the germ-free recipients (n = 3-5). After one month of colonization, tissues were collected from the inoculated ex-germ-free mice and stored at −80 °C until further analysis.

### Total RNA extraction and RT-qPCR

Total RNA was isolated from the large intestine using RNA-Bee (Tel-Test Inc., Friendswood, TX), as previously described (Li *et al*., 2017). RNA concentrations were quantified using a NanoDrop 1000 Spectrophotometer (Thermo Scientific, Waltham, MA) at 260 nm. The integrity of total RNA samples was evaluated by formaldehyde-agarose gel electrophoresis with visualization of 18S and 28S rRNA bands under UV light. Extracted RNA samples were then reverse transcribed to cDNA using a High-Capacity cDNA Reverse Transcription Kit (Life Technologies, CA). The resulting cDNA products were amplified by qPCR, using a Sso Advanced Universal SYBR Green Supermix in a Bio-Rad CFX384 Real-Time PCR Detection System (Bio-Rad, Hercules, CA). Data were normalized to the housekeeping gene Glyceraldehyde 3-phosphate dehydrogenase (Gapdh) using the ΔΔCq method and were expressed as % of Gapdh. Primer sequences are shown in Supplementary Table 7.

## RESULTS

### Investigation of cell type-specific responses following neonatal exposure to BDE-99 in late adulthood

To investigate the transcriptomic signatures in late adulthood as regulated by neonatal BDE-99 exposure (from postnatal day [PND] 2-4), we performed scRNA-seq in livers of 15-month-old male adults. ScRNA-seq was able to identify the major hepatic resident cells (i.e. hepatocytes, cholangiocytes, endothelial cells, stellate cells, myofibroblasts, Kupffer cells), as well as immune cells that circulate into liver via blood (i.e. B cells, T cell populations, CD4 T cells, CD8 T cells, natural killer [NK] cells, conventional dendritic cells [cDC], plasmacytoid DCs [pDC], neutrophils, and basophils) (Fig. 2A). We observed minimal difference in the overall clustering and cell type labeling between vehicle- and BDE-99-exposed groups (Fig. S1). Cell types in the single cell data clusters were identified and labeled using established unique marker genes (as summarized in Table S1) for each liver cell type (Guilliams *et al*., 2022). As expected, each liver cell type expressed the corresponding unique marker gene(s) (Fig. 2B).

**Figure 2.**
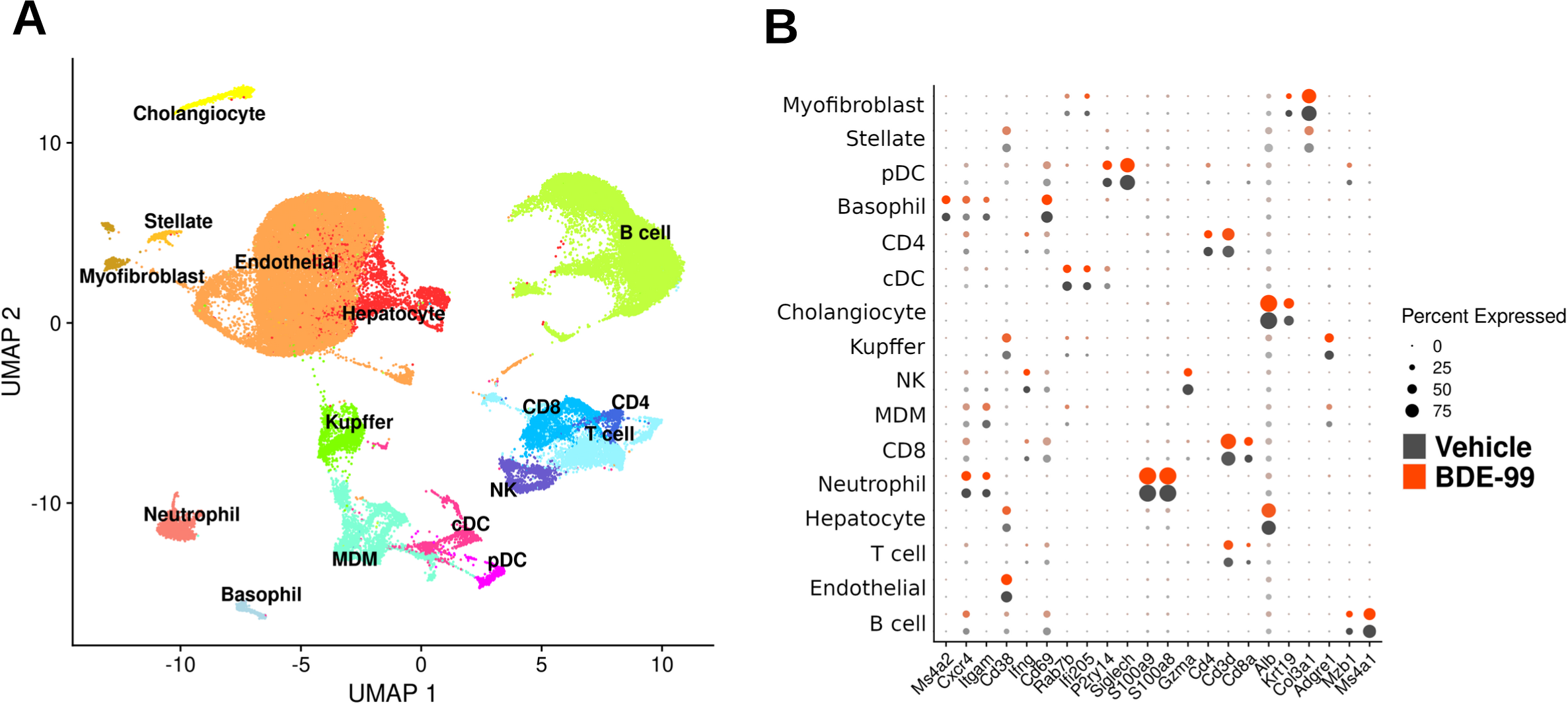
Clustering and visualization of specific marker genes in each cell type of liver. **A.** Visualization of all labeled cell clusters detected through scRNA-seq (vehicle and BDE-99 exposed groups were combined). The first two uniform manifold approximation and projections (UMAP) were used. **B.** Representation of key marker genes used for cell type labeling. Each marker gene was uniquely enriched for the corresponding cell type cluster in both vehicle and BDE-99-exposed groups. Black and red colors indicate vehicle and BDE-99 exposed groups, respectively.

To compare the metabolic capacity of liver cell types, we performed gene ontology enrichment of all expressed genes related to xenobiotic biotransformation (Fig. S2). Genes enriched in hepatocytes are mainly involved in the metabolism of fatty acids, retinoids, steroids, bile acids, xenobiotics, and hormones. Endothelial cells and stellate cells were enriched in genes involved in response to toxic substances. Myofibroblasts were enriched in the metabolism of hormones, retinoids, xenobiotics, and fatty acids. Basophils had an enrichment in genes involved in lipoxygenase pathway and icosanoid metabolism. Kupffer cells and other immune cells (e.g. MDM, neutrophil) had an enrichment in genes involved in the cell redox cycle, as well as icosanoid and lipoxygenase metabolism.

### Persistent down-regulation of major drug-processing genes following early life BDE-99 exposure

One of the critical functions of the liver is detoxification through drug-processing genes, including phase-I and -II drug metabolizing enzymes and transporters (Grant, 1991; Cui *et al*., 2009; Parkinson *et al*.). At PND 60, using bulk RNA-Seq, we recently showed that early life exposure to BDE-99 resulted in a persistent down-regulation of these drug-processing genes in liver of young adults, with males being more susceptible than females (Lim *et al*., 2021). As a follow-up to our previous study, we investigated the extent of early life BDE-99 on the reprogramming of drug-processing genes in liver of male offspring in late adulthood at single cell resolution.

Neonatal BDE-99 exposure resulted in persistently dysregulated distinct drug-processing genes in various cell types of livers of 15-month-old adults (Fig. 3 and Table S2). In hepatocytes, overall, most of the differentially regulated drug-processing genes were down-regulated by neonatal BDE-99 exposure (Fig. 3A). For example, neonatal BDE-99 exposure persistently down-regulated cytochrome P450 (Cyp) *1a2*, *Cyp3a11*, and *Cyp4a10*, which are the prototypical target genes for the major xenobiotic-sensing transcription factors aryl hydrocarbon receptor (Ahr), pregnane X receptor (PXR), and peroxisome proliferator-activated receptor alpha (PPARα) respectively, in hepatocytes of late adulthood. In general, neonatal BDE-99 exposure down-regulated the families of aldehyde dehydrogenase (*Aldh*), carboxylesterase (*Ces*), *Cyp*s, glutathione-*S* transferase (*Gst*), UDP-glucuronosyltransferase (*Ugt*) were down-regulated in hepatocytes in late adulthood. Regarding transporters, with the exception of ATP binding cassette c9 (*Abcc9*/multidrug resistance-related protein 9 [Mrp9]) and solute carrier 47a1 (*Slc47a1*) (Multidrug and toxin extrusion protein 1, Mate1), several transporters important for xenobiotic disposition and bile flow were persistently down-regulated by neonatal BDE-99 exposure in hepatocytes. These down-regulated transporters included organic anion transporting polypeptide (Oatp1a1/*Slco1a1*), *Abcc2* (multidrug resistance-associated protein 2, Mrp2), *Abcc3* (Mrp3), *Abcb1b* (P-glycoprotein), *Abcb4* (multidrug resistance protein 3, Mdr2), *Slc10a1* (sodium taurocholic cotransporting polypeptide, Ntcp), *Slc22a1* (organic cation transporter, Oct1), *Slc22a7* (organic anion transporter 2, Oat2), and *Slc27a5* (bile acyl-CoA synthetase).

**Figure 3.**
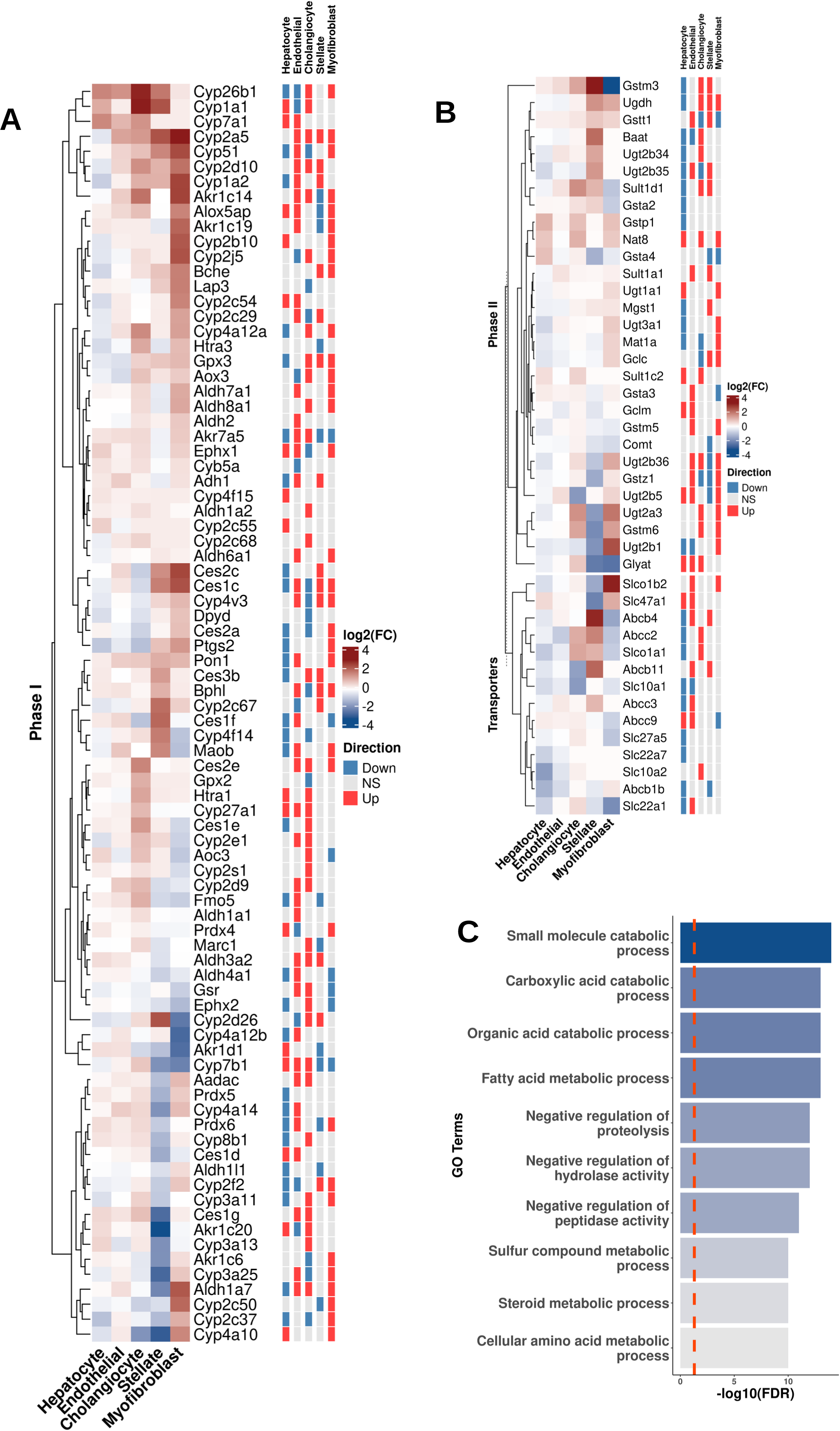
Persistently dysregulated expression signatures of drug-processing genes in resident hepatic cell populations at 15 months of age following neonatal exposure to BDE-99. **A.** Phase-I drug-processing enzymes that were differentially expressed (rows) in at least one of the major hepatic resident cell types (columns) with xenobiotic biotransformation capabilities are shown in a heatmap (left side). The colors of the heatmap represent the log_2_ fold change of liver genes of the BDE-99 exposed adults as compared to the vehicle control. “Direction” indicates whether a gene is up- or down-regulated from neonatal exposure to BDE-99 (Bonferroni-adjusted *p*-value < 0.05). **B.** Phase-II enzymes and transporters involved in xenobiotic metabolism processes that were differentially expressed (rows) in at least one of the major hepatic resident cell types (columns) with xenobiotic biotransformation capabilities are shown in a heatmap (left side). The colors of the heatmap represent the log_2_ fold change of liver genes of the BDE-99 exposed adults as compared to the vehicle control. “Direction” indicates whether a gene is up- or down-regulated from neonatal exposure to BDE-99 (Bonferroni-adjusted *p*-value < 0.05). **C.** Top 10 Gene Ontology enrichment of down-regulated genes in hepatocytes following neonatal exposure to BDE-99.

Interestingly, in addition to hepatocytes, neonatal BDE-99 exposure also persistently dysregulated distinct drug-processing genes in non-parenchymal cells (i.e. endothelial cells, cholangiocytes, stellate cells, and myofibroblasts) in livers of adults. Contrary to a general decreasing trend of the expression of drug-processing genes in hepatocytes, most of the BDE-99 regulated drug-processing genes in non-parenchymal cells were up-regulated (Fig. 3). Correspondingly, several gene ontology enrichment terms related to xenobiotic and fatty acid metabolism in the non-parenchymal cell types were also persistently up-regulated (e.g. fatty acid metabolic process, response to xenobiotic stimulus; Fig. S3A). Specifically, families of *Aldh*, *Ces*, *Cyp*, *Gst*, *Ugt*, as well as several transporters (e.g. Mdr2 in endothelial cells and stellate cells; Mrp2 in cholangiocytes; Mrp3, Oct1, and Mate1 in endothelial cells; Oatp1b2 in endothelial cells and myofibroblasts) were up-regulated in non-parenchymal cell types. As noted above, these genes were down-regulated in hepatocytes (Fig. 3A, Fig. 3B, and Table S2-S3). The up-regulation of distinct drug-processing genes in non-parenchymal cell types may represent a compensatory mechanism for chemical detoxification due to BDE-99 mediated persistent decrease in their expression in hepatocytes. However, it should be noted that the fold change of expression levels of the drug-processing genes in non-parenchymal cells was much lower than those in hepatocytes under both basal conditions and following early life BDE-99 exposure (Fig. S4, Table S4-S5). Thus, the persistent BDE-99 mediated up-regulation of drug-processing genes in non-parenchymal cells may not sufficiently compensate the down-regulated expression of drug-processing genes in hepatocytes (Fig. 3C). In summary, neonatal BDE-99 exposure persistently dysregulated distinct drug-processing genes not only in young adulthood as we reported previously (Lim *et al*., 2021), but also in late adulthood; in addition, scRNA-Seq revealed opposite regulatory patterns of drug-processing genes by neonatal BDE-99 exposure in hepatocytes (down-regulation), as compared to non-parenchymal cells (up-regulation).

### Neonatal BDE-99 exposure persistently increased the hepatic proportion of immune cells in liver in late adulthood

As we previously reported using bulk RNA-Seq, neonatal BDE-99 exposure produced a pro-inflammatory transcriptomic signature in adult male livers at PND 60 (Lim *et al*., 2021). To determine at single cell resolution to what extent such regulatory pattern persists into late adulthood, and how various cell types in liver contribute to inflammation, we investigated the changes in cell type proportions in late adulthood following neonatal BDE-99 exposure. Interestingly, there was an increasing trend in the overall proportions of circulating immune cell types in liver following neonatal BDE-99 exposure. Among these immune cell types, there was a statistically significant increase in the neutrophil proportions by neonatal BDE-99 exposure (*p* < 0.05) (Fig. 4A). In addition, although not statistically significant, the proportions of T cell, MDM, NK, cDC, basophil, and myofibroblast populations all tended to increase by neonatal BDE-99 exposure.

**Figure 4.**
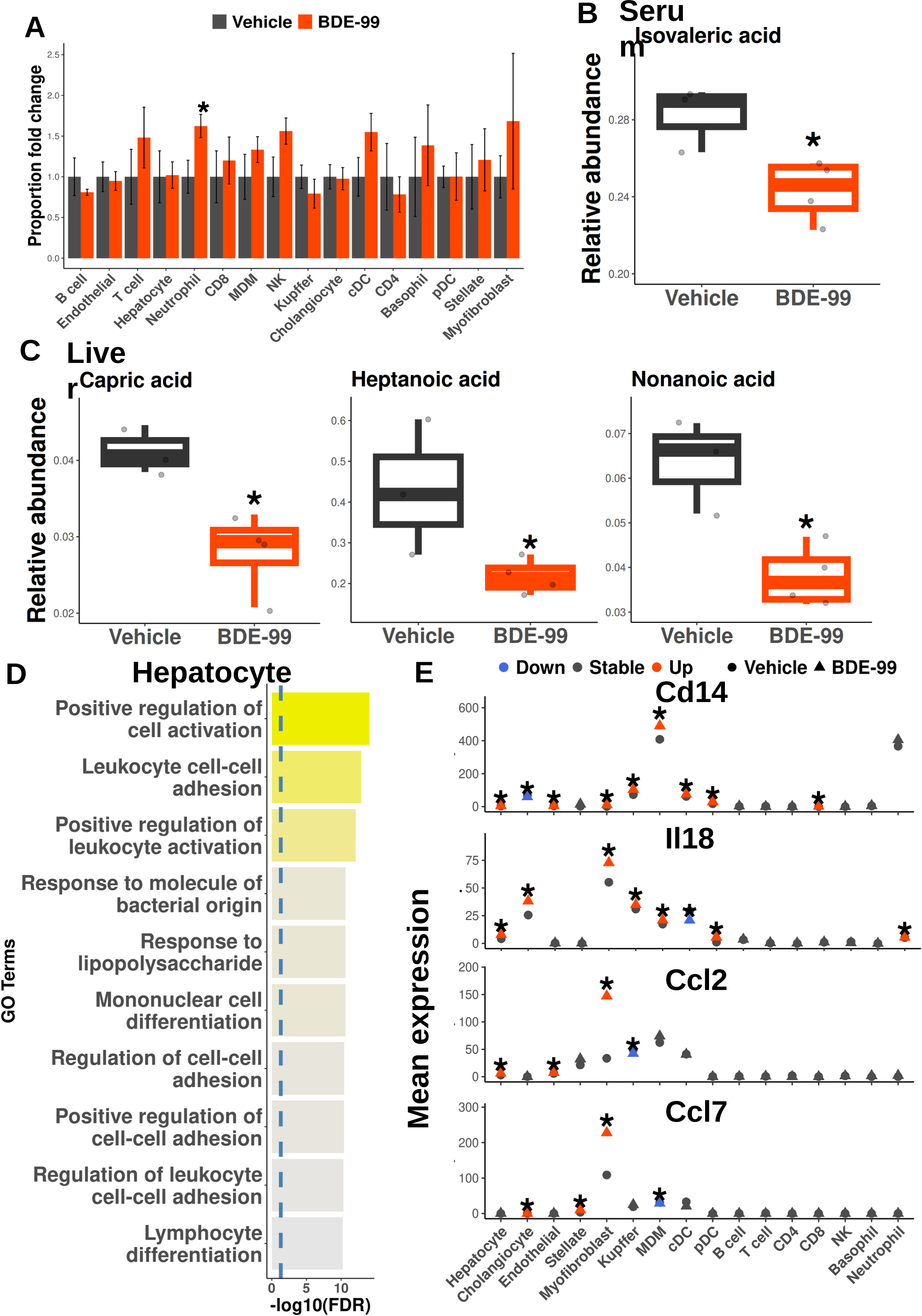
**A.** Changes in proportions of the liver cell types in 15-month old adult mouse livers following neonatal exposure to BDE-99. Y-axis shows the ratio of the % of each cell type of BDE-99 exposed group over that of the vehicle-exposed group. Asterisks represent *p*-value < 0.05 (two-way t-test with assumption of unequal variance). **B.** Relative abundance of isovaleric acid detected in serum. Asterisks represent *p*-value < 0.05 (one way ANOVA with Tukey’s post-hoc test). **C.** Relative abundance of medium-chain fatty acids detected in liver. Asterisks represent *p*-value < 0.05 (one way ANOVA with Tukey’s post-hoc test). **D.** Top 10 up-regulated gene ontology terms in hepatocytes from 15-month-old males by early life exposure to BDE-99. **E.** Average expression of pro-inflammatory markers in hepatic cell types following neonatal exposure to BDE-99. Red and blue colors represent up- and down-regulation, respectively. Vehicle and BDE-99-exposed groups are shown as circles and triangles, respectively. Asterisks indicate differential expression (Bonferroni-adjusted *p*-value < 0.05).

The neonatal BDE-99 mediated increase in hepatic immune cell proportions is consistent with a decrease in several anti-inflammatory fatty in serum and liver in late adulthood. For example, neonatal BDE-99 exposure decreased isovaleric acid in serum (Fig. 4B). Isovaleric acid is linked to anti-inflammatory processes (Nakkarach *et al*., 2021) and the decrease of its relative expression suggests a potential systemic inflammatory environment. Neonatal BDE-99 exposure also persistently decreased various medium-chain fatty acids in the liver, including capric acid, heptanoic acid, and nonanoic acid (Fig. 4C). Medium-chain fatty acids are associated with anti-inflammatory properties (Jia *et al*., 2020; Rial *et al*., 2016).

In late adulthood, the neonatal BDE-99 mediated increase in hepatic immune cell proportions is consistent with a pro-inflammatory transcriptomic signature in hepatocytes, including an up-regulation in genes involved in leukocyte cell-cell adhesion and activation, mononuclear cell differentiation, and lymphocyte differentiation (Fig. 4D). In addition to pro-inflammation, genes involved in response to molecule of bacterial origin and lipopolysaccharide were up-regulated in hepatocytes by neonatal BDE-99 exposure (Fig. 4D), suggesting that the pro-inflammatory state in liver may be originated from a dysregulated gut environment.

Specific examples of persistently regulated cytokines/chemokines by neonatal BDE-99 exposure are shown in Fig. 4E. Cluster of differentiation 14 (*Cd14*), which is a part of the pathogen-associated molecular pattern (PAMP) detection system, was expressed the highest in MDM, and neonatal BDE-99 exposure persistently up-regulated its expression in hepatocytes, endothelial cells, myofibroblasts, Kupffer cells, MDM, cDC, pDC, and CD8 T cells, and was down-regulated in cholangiocytes (Fig. 4E). The pro-inflammatory cytokine interleukin 18 (*Il18*) was expressed the highest in myofibroblasts, and was persistently up-regulated by neonatal BDE-99 exposure in resident liver cells (i.e. hepatocytes, cholangiocytes, myofibroblasts, Kupffer cells), as well as MDM, pDC, and neutrophils, and was down-regulated in cDC (Fig. 4E). CC motif chemokine ligand (*Ccl*)*2* and *7* are involved in macrophage and neutrophil recruitment (Gschwandtner *et al*., 2019; Xie *et al*., 2021), and both of them were expressed the highest in myofibroblasts (Fig. 4E). *Ccl2* was persistently up-regulated by neonatal BDE-99 exposure in hepatocytes, endothelial cells, and myofibroblasts, and was down-regulated in Kupffer cells. *Ccl7* was persistently up-regulated by neonatal BDE-99 exposure in cholangiocytes, stellate cells, and myofibroblasts, and was down-regulated in MDM (Fig. 4E).

### Increased macrophage-centered pro-inflammatory signaling by neonatal BDE-99 exposure

The persistent up-regulation of markers involved in inflammation and immune cell recruitment may indicate enhanced interactions between hepatocytes and non-parenchymal cells to promote a pro-inflammatory state. Therefore, to investigate the regulation of the cell-cell interactive networks by neonatal BDE-99 exposure, we conducted ligand-receptor mediated intercellular communication analysis centering on hepatic inflammation (Fig. 5). The macrophage migration inhibitory factor (MIF) acts as a cytokine that turns on pro-inflammatory responses in macrophage-related cells (Roger *et al*., 2001; Calandra and Roger, 2003). MIF signaling is up-regulated upon detection of bacterial antigens and as a response to PAMPs (Roger *et al*., 2001; Calandra and Roger, 2003). Intercellular signaling analysis showed that neonatal BDE-99 exposure persistently up-regulated MIF signaling in macrophage-related hepatic immune cell populations (i.e. MDM, Kupffer cell, cDC, and pDC). The increased MIF signal is predicted to be sent from cell types including hepatocytes, myofibroblasts, and pDCs (Fig. 5A). In addition, genes related to leukocyte migration and chemotaxis were persistently up-regulated in Kupffer cells and MDMs by neonatal BDE-99 exposure (Fig. 5B). Similarly, chemotaxis signatures and cytotoxic response-related genes (i.e. cell killing, adaptive immune response) were also up-regulated in cDC and pDC populations following neonatal BDE-99 exposure (Fig. S5A). Interestingly, fatty acid metabolism-related signatures were also persistently up-regulated following early life exposure to BDE-99 (Fig. 6B). To note, increased lipid metabolism in macrophages plays a critical role in their activation (Remmerie and Scott, 2018; Batista-Gonzalez *et al*., 2019). In the leukocyte populations (Kupffer cells, MDM, cDC, and pDC), leukocyte migration and inflammatory response were persistently down-regulated (Fig. S5B and Fig. S6), suggesting an overall up-regulation of migratory and pro-inflammatory responses following neonatal exposure to BDE-99. Gene expression signatures related to cell chemotaxis and leukocyte migration were persistently up-regulated in myofibroblasts, and tissue remodeling and wounding responses were persistently down-regulated by neonatal exposure to BDE-99 (Fig. S7A and Fig. S7B). Concordant to the predicted up-regulation of MIF signaling, neonatal BDE-99 exposure persistently increased the number of Kupffer cells and the Kupffer cell specific expression of the pro-inflammatory chemokine *Cxcl10*. Neonatal BDE-99 exposure also persistently increased the number of MDM and the MDM-specific expression of the pro-inflammatory cytokines Cxcl10 and Il6 (Fig. 5C). These results indicate that the activation of hepatic macrophage-related cells is important in promoting the persistent inflammatory signatures following neonatal BDE-99 exposure.

**Figure 5.**
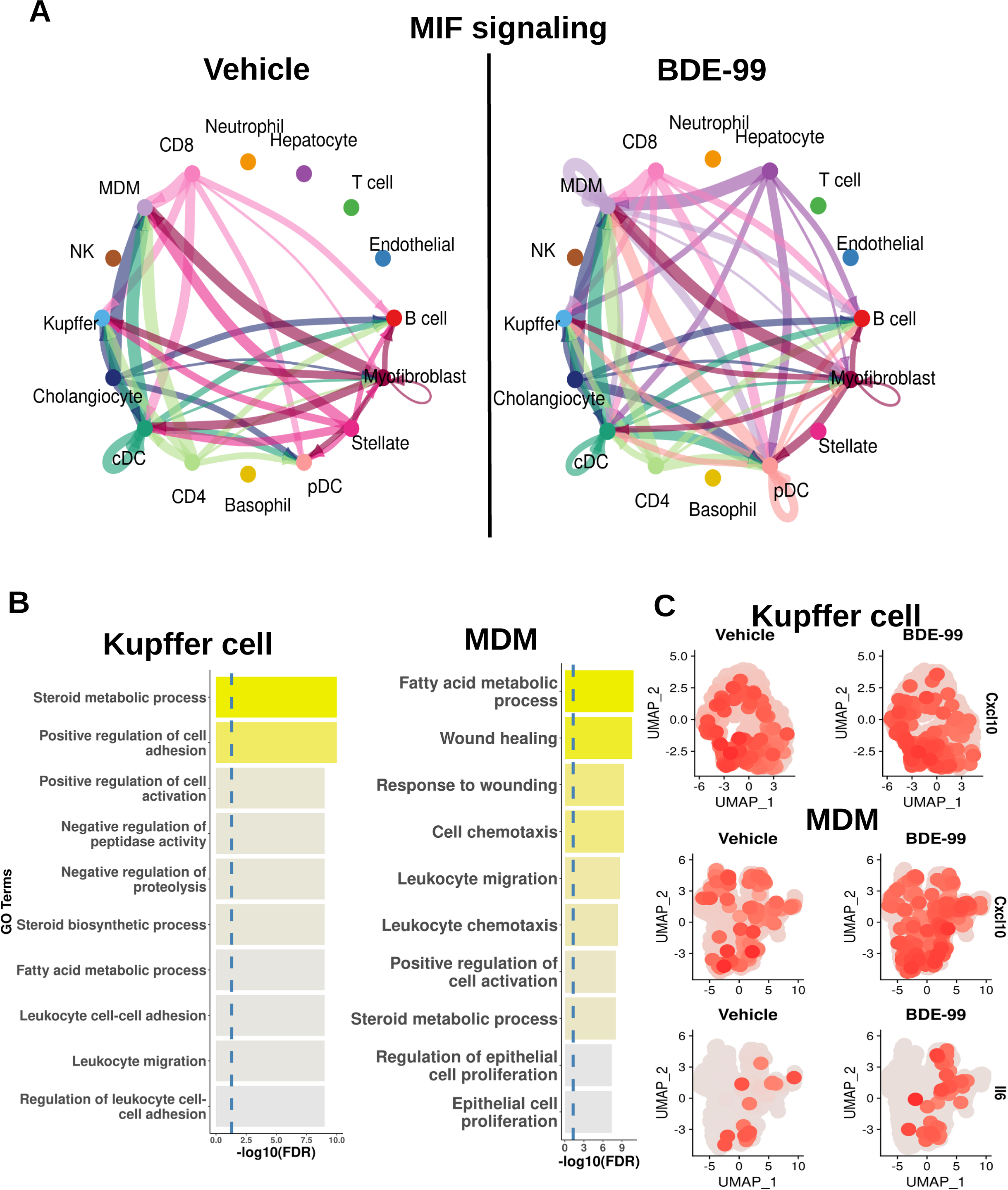
Increased macrophage migration inhibitory factor (MIF) signaling among multiple liver cell types and changes in downstream immune signatures in adult mouse livers following early life exposure to BDE-99. **A.** Visualization of the cell-cell communications of the MIF signaling pathway in in vehicle and BDE-99-exposed groups. Each cell type contains a unique color, and the matched colors represent signal communication direction from one cell type to another. The thickness of the arrows represents the probability of communication. **B.** Top 10 up-regulated gene ontology enrichment results in Kupffer cells and MDMs in adult mouse livers following neonatal exposure to BDE-99. **C.** Up-regulated expression of pro-inflammatory markers in Kupffer cells and MDMs. Grey and red indicate low and high expression, respectively.

**Figure 6.**
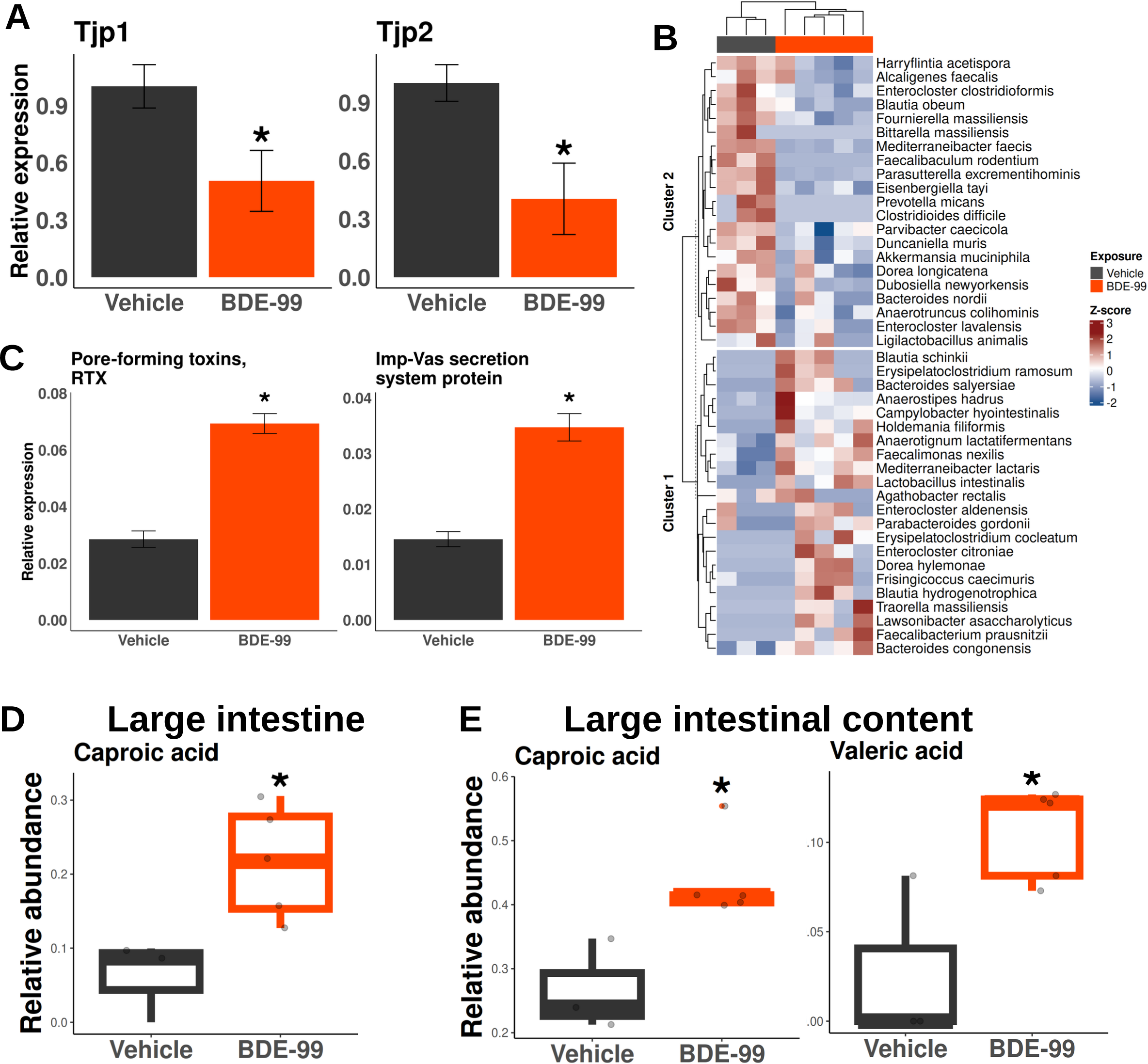
Evidence of dysregulated gut environment by early life exposure to BDE-99. **A.** Down-regulated expression of tight junction proteins by RT-qPCR in the large intestine in 15-month-old males following neonatal exposure to BDE-99. Asterisks show *p*-value < 0.05 (two-way t test with assumption of unequal variance). **B.** Persistent dysbiosis by early life exposure to BDE-99 at the full species level. Two main clusters are formed by k-means (k = 2). Top bars represent adults neonatally exposed to vehicle (black) or BDE-99 (orange). Red and blue colors show statistically significant increase or decrease of taxa abundance, respectively (ANCOM-BC, BH-adjusted *p*-value < 0.05). **C.** Persistent up-regulation of predicted functions linked to host cell damage from dysbiotic gut. Persistent increase of metabolites in the large intestinal cell (**D**) and large intestinal content (**E**).

To seek phenotypic evidence of hepatic inflammation and related disease, we conducted H&E staining of the liver (Fig. S8A). Overall, mild hepatic inflammation (Fig. S8A) and bile duct hyperplasia (Fig. S8A) were observed following neonatal exposure to BDE-99 (Fig. S8B). The gross phenotypic changes in liver histology may be contributed by neonatal BDE-99 exposure mediated increase in hepatic proportions of circulating immune cells and myofibroblasts as discussed above.

### Altered gut environment as a potential mediator of hepatic pro-inflammation

One mechanism of persistent hepatic inflammation and increased influx of circulating immune cells is dysregulated gut environment (Tripathi *et al*., 2018; Phipps *et al*., 2020; Barrow *et al*., 2021). The large intestine harbors the most diverse gut microflora (Hillman *et al*., 2017), and decreased intestinal barrier is closely linked to dysregulation of the gut microbiota (Kinashi and Hase, 2021; Stolfi *et al*., 2022). As mentioned above, associated with pro-inflammation, neonatal BDE-99 exposure persistently increased the expression of genes involved in response to bacterial components (Fig. 4C), suggesting that the pro-inflammatory state in liver may be originated from a dysregulated gut environment. Therefore, we hypothesize that neonatal BDE-99 exposure disrupted gut barrier functions and produced a dysbiosis of the gut microbiome in late adulthood.

The mRNA expression of tight junction proteins (Tjp1 and 2) in the large intestine at 15 months of age were both persistently down-regulated by neonatal BDE-99 early life exposure (Fig. 6A). These results suggest that neonatal exposure to BDE-99 may increase gut permeability. We previously reported that early life exposure to BDE-99 reprogrammed the dysregulated key taxa linked to liver diseases in the large intestine at PND 60 (Lim *et al*., 2021). At 15 months post-exposure to BDE-99, we determined whether there was a long-term dysbiosis of the large intestinal flora. Metagenomic shotgun sequencing of the large intestinal content showed 43 differentially abundant taxa at the full species level (Fig. 6B). Neonatal BDE-99 exposure persistently down-regulated *Harryflintia acetispora*, *Alcaligenes faecalis*, *Enterocloster clostridioformis*, *Blautia obeum*, *Fournierella massiliensis*, *Bittarella massiliensis*, *Mediterraneibacter faecis*, *Faecalibacculum rodentium*, *Parasutterella excrementihominis*, *Eisenbergiella tayi*, *Prevotella micans*, *Clostridioides difficile*, *Parvibacter caecicola*, *Duncaniella muris*, *Akkermansia muciniphila*, *Dorea longicatena*, *Dubosiella newyorkensis*, *Bacteroides nordii*, *Anaerotruncus colihominis*, *Enterocloster lavalensis*, and *Ligilactobacillus animalis*. Conversely, neonatal BDE-99 exposure persistently up-regulated *Blautia schinkii*, *Erysipelatoclostridium ramosum*, *Bacteroides salyersiae*, *Anaerostipes hadrus*, *Campylobacter hyointestinalis*, *Holdemania filiformis*, *Anaerotignum lactatifermentans*, *Faecalimonas nexilis*, *Mediterraneibacter lactaris*, *Lactobacillus intestinalis*, *Agathobacter rectalis*, *Enterocloster aldenensis*, *Parabacteroides gordonii*, *Erysipelatoclostridium cocleatum*, *Enterocloster citroniae*, *Dorea hylemonae*, *Frisingicoccus caecimuris*, *Blautia hydrogenotrophica*, *Traorella massiliensis*, *Lawsonibacter asaccharolyticus*, *Faecalibacterium prausnitzii*, *Bacteroides congolensis*. Functional predictions showed that the up-regulated taxa were associated with cytotoxic secretion that damages host cells (Fig. 6C). Both in the host large intestinal tissue and large intestinal content, the relative abundance of the inflammation-associated caproic acid (Nakkarach *et al*., 2021) was persistently up-regulated by neonatal BDE-99 exposure (Fig. 6D-6E). Valeric acid, which has been shown to be associated with both pro- and anti-inflammatory processes (Nakkarach *et al*., 2021; Lai *et al*., 2021), was also up-regulated by neonatal BDE-99 exposure to BDE-99 (Fig. 6E). Coupled with dysregulated microbiota and their constituents, these may aggravate the hepatic pro-inflammation and microbial component influx following neonatal BDE-99 exposure.

In summary, our data showed that neonatal BDE-99 exposure persistently altered the hepatic transcriptomic signatures in a cell-type-specific manner in late adulthood; while the major drug-processing genes were persistently down-regulated in hepatocytes, they were up-regulated in non-parenchymal cells. In addition, neonatal BDE-99 exposure persistently increased hepatic infiltration of immune cells and expression of pro-inflammatory cytokines/chemokines, as well as enhanced multi-cellular communications to promote inflammation. Such a pro-inflammatory state in the liver due to neonatal BDE-99 exposure is associated with reduced intestinal tight junction protein expression, increased abundance of harmful microbes in the gut, as well as a pro-inflammatory signature of fatty acid metabolites within the gut-liver axis. Because microbial LPS signaling is known to induce pro-inflammatory liver injuries (Hamesch *et al*., 2015), we hypothesized that neonatal BDE-99 exposure-mediated hepatic pro-inflammatory gene expression signatures may at least partially be contributed by the dysregulated gut microbiome and gut environment.

As a first step to determine the necessity of gut microbiome in hepatic immune response in a cell-type specific manner, we used scRNA-Seq to determine the basal functions of the gut microbiome in hepatic immune pathways using age-matched conventional and germ-free adult mice. Because immune signaling involves cell-cell interactions, we conducted ligand-receptor-mediated intercellular communication analysis. In livers of control conventional mice, the cellular composition is similar to the control mice from the BDE-99 exposure study (Fig. 7A and Fig. S9). In livers of the germ-free mice, the absence of the gut microbiome up-regulated the pro-inflammatory complement signaling involving Kupffer cells and MDMs as targeted by hepatocyte, cholangiocyte, and myofibroblast populations (Fig. 7B). Cell chemotaxis, major histocompatibility complex II assembly, and pro-inflammation-related signatures were up-regulated in the absence of microbiome in hepatocytes (Fig. S10A and Table S6). MHC class II-expressing hepatocytes are found in clinical hepatitis (Mehrfeld *et al*., 2018; Lu *et al*., 2020; Herkel *et al*., 2003), suggesting a role of the microbiome in hepatic immune regulation. Response to wounding-related signatures was up-regulated in cholangiocytes and immune cell migration, and immune effector process-related signatures were up-regulated in myofibroblasts in the absence of microbiome (Fig. S10A). The absence of microbiome resulted in up-regulated signatures involved in wounding responses in both of the predicted target cells of complement signaling (i.e. Kupffer cells and MDMs), with Kupffer cells showing up-regulated adaptive immune response-related signatures (Fig. S10B). Furthermore, the mRNAs of several pro-inflammatory markers and regulators were up-regulated in other hepatic cell types in the absence of microbiome (Fig. 7C). Specifically, tumor necrosis factor-alpha (*Tnfα*) was up-regulated in Kupffer cell, MDM, cDC, T cell, basophil, and neutrophil populations, while its expression was down-regulated in cholangiocytes, pDC, γδT cell populations in the absence of microbiome. Vascular endothelial growth factor A (*Vegfa*) was up-regulated in all hepatic cell types except stellate cell and Kupffer cell populations (down-regulated in both cell types) and B cell, B1 cell, and T cell populations in the absence of microbiome. Early growth response 1 (*Egr1*) was up-regulated in endothelial cells, stellate cells, Kupffer cells, MDMs, T cells, and neutrophils, while down-regulated in pDC, B cell, and NK cell populations in the absence of microbiome. *Cxcl13* was up-regulated in all detected hepatic cell types except hepatocytes (down-regulated), pDC, B cell, B1 cell, γδT cell, basophil, and neutrophil populations in the absence of microbiome. The intercellular adhesion molecule 1 (*Icam1*) was up-regulated in endothelial cells, stellate cells, MDMs, pDCs, NK cells, and neutrophils but was down-regulated in hepatocytes, B1 cells, and basophils in GF conditions. *Cxcl12* was up-regulated in endothelial cells, myofibroblasts, Kupffer cells, cDCs, and T cells, and was down-regulated in hepatocytes, cholangiocytes, and B cells in the absence of microbiome. *Il1α* was up-regulated in cholangiocytes, endothelial cells, myofibroblasts, Kupffer cells, and neutrophils, though down-regulated in hepatocytes, cDC, B cell, B1 cell, and T cell populations in GF livers. *Il5* was up-regulated in MDM and T cells in the absence of microbiome. Overall, the absence of microbiome showed consistent signatures of increased hepatic pro-inflammation. These aberrant transcriptomic signatures at single cell resolution indicate that the presence of a normal microbiome is necessary for maintaining hepatic immunotolerance and highlights the importance of the gut-liver axis in regulating hepatic immune functions.

**Figure 7.**
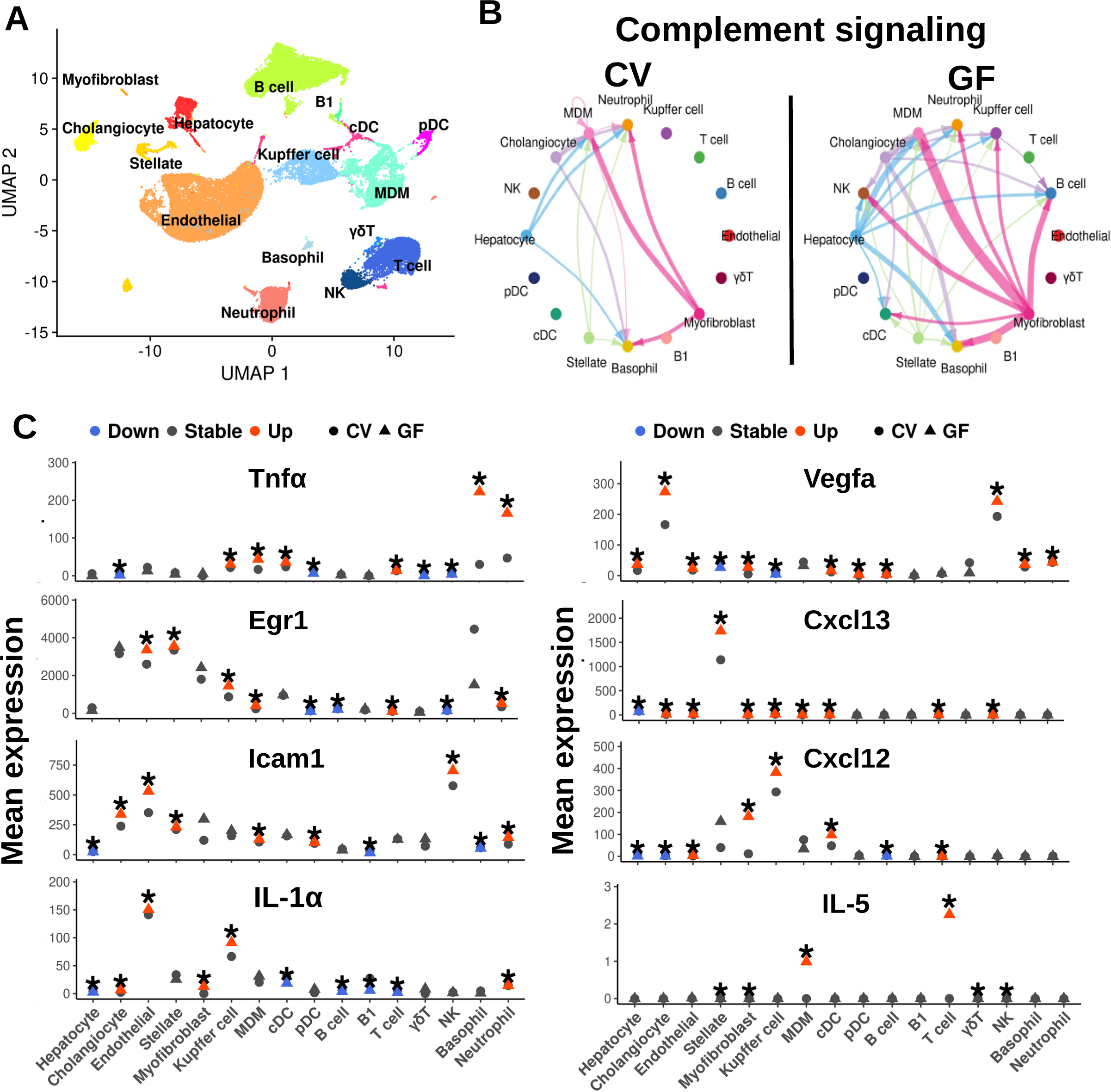
Up-regulated immune hepatic signaling patterns in the absence of gut microbiome. **A.** UMAP representation of cell clusters labeled by cell type in both CV and GF mice. **B.** Cell-cell communication signaling visualization for the complement signaling pathway in CV and GF livers. Each cell type contains a unique color and the matched colors represent signal communication direction from one cell type to another. The thickness of the arrows shows the probability of communication. **C.** Average expression of pro-inflammatory markers and immune regulators in CV and GF liver cell types. Red and blue colors represent up- and down-regulation, respectively. Vehicle and BDE-99-exposed groups are shown as circles and triangles, respectively. Asterisks indicate differential expression (Bonferroni-adjusted *p*-value < 0.05).

### Transplantation of dysregulated microbiome promotes a pro-inflammatory gut environment

To further link the functional capacity of the altered microbial composition by early life exposure to BDE-99, we transplanted the large intestinal content of the adults that were neonatally exposed to vehicle or BDE-99 to germ-free adult mice (Fig. 8A). Upon transplantation, mRNA levels of tight junction proteins were down-regulated in ex germ-free mice transplanted with gut microbiome from the BDE-99 donors (Fig. 8B). One important regulator of gut barrier maintenance is by interleukin (*Il*) *22* (Keir *et al*., 2020). The expression of *Il22* in BDE-99 donors tended to decrease (Fig. 8B). *Il17*, which is linked to pro-inflammatory processes (Mills, 2023), and *TNFα* were up-regulated by the transplantation of BDE-99 large intestinal content (Fig. 8B). These results suggest that the altered gut microbiota in the large intestine can act as initiators of inflammatory processes in the gut environment and may further impact the pro-inflammatory events that were observed by early life exposure to BDE-99.

**Figure 8.**
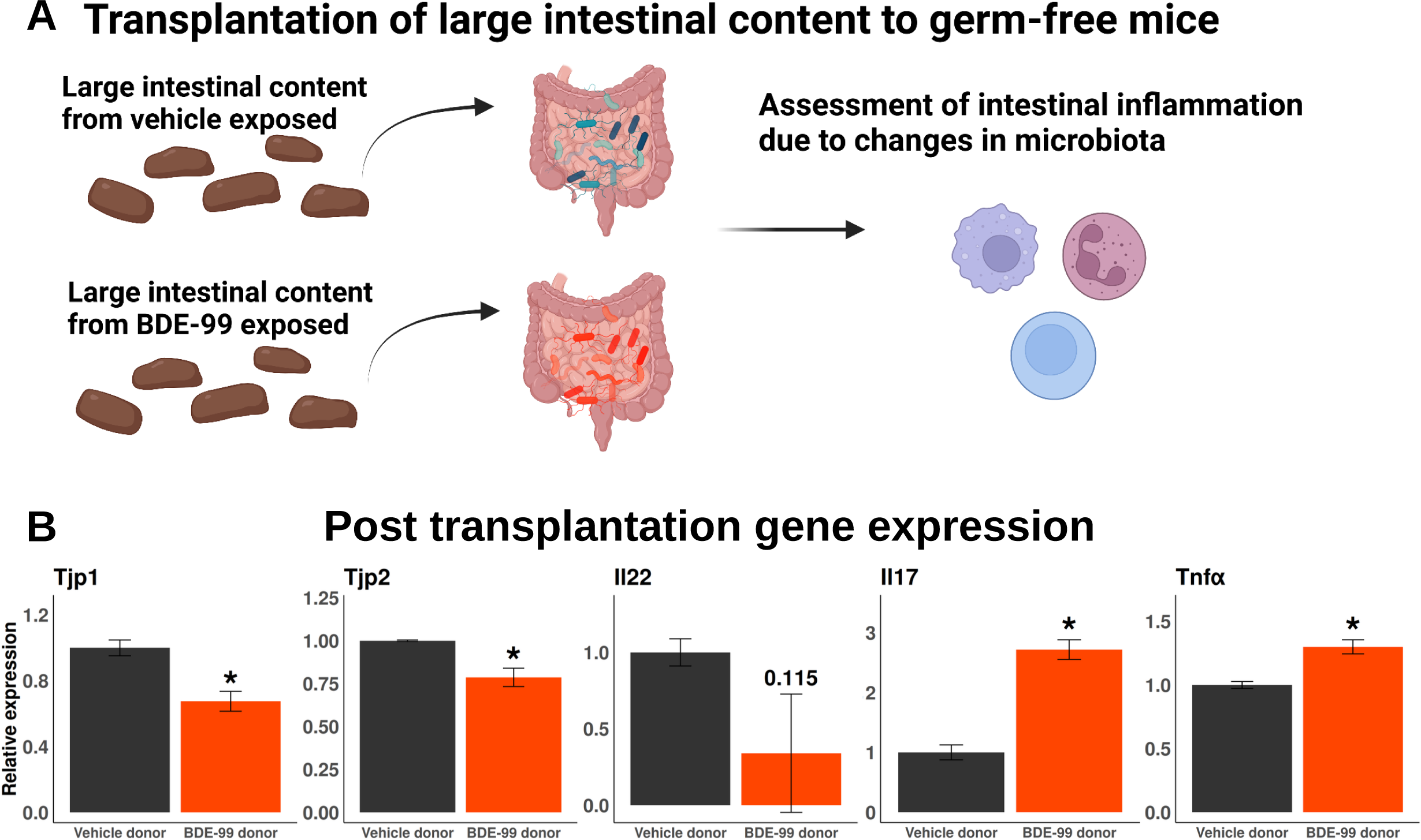
Changes in microbiome composition by early life exposure to BDE-99 is linked to regulation of inflammation. **A.** Experimental schematic of fecal microbiota transplantation to adult male germ-free mice. To study the impact on the immuno-modulation in the gut environment from the changes in the microbiome from early life exposure to BDE-99, large intestinal content was transplanted to adult germ-free mice. **B.** Down-regulation of the transcripts of tight junction proteins and up-regulated pro-inflammatory cytokines in the large intestine post-transplantation of large intestinal content from adults neonatally exposed to vehicle or BDE-99 in germ-free mice, as determined by RT-qPCR. Asterisks represent *p* < 0.05 (two-way t-test with unequal variance).

## DISCUSSION

In the present study, we investigated the long-term hepatotoxic effect and the underlying molecular mechanisms in late adulthood in males following neonatal BDE-99 exposure. Using single cell transcriptomics, we were able to achieve higher precision and resolution as compared to bulk RNA-Seq method to address the cell-cell interactive networks during the adult onset of hepatotoxicity following early life PBDE exposure (Fig. 9). Using germ-free mice with or without fecal microbiome transport from BDE-99 exposed conventional mice, we demonstrated the necessity of gut microbiome in maintaining the immunotolerance in liver, and the mechanistic involvement of the “diseased” microbiome in PBDE-mediated hepatic reprogramming.

**Figure 9.**
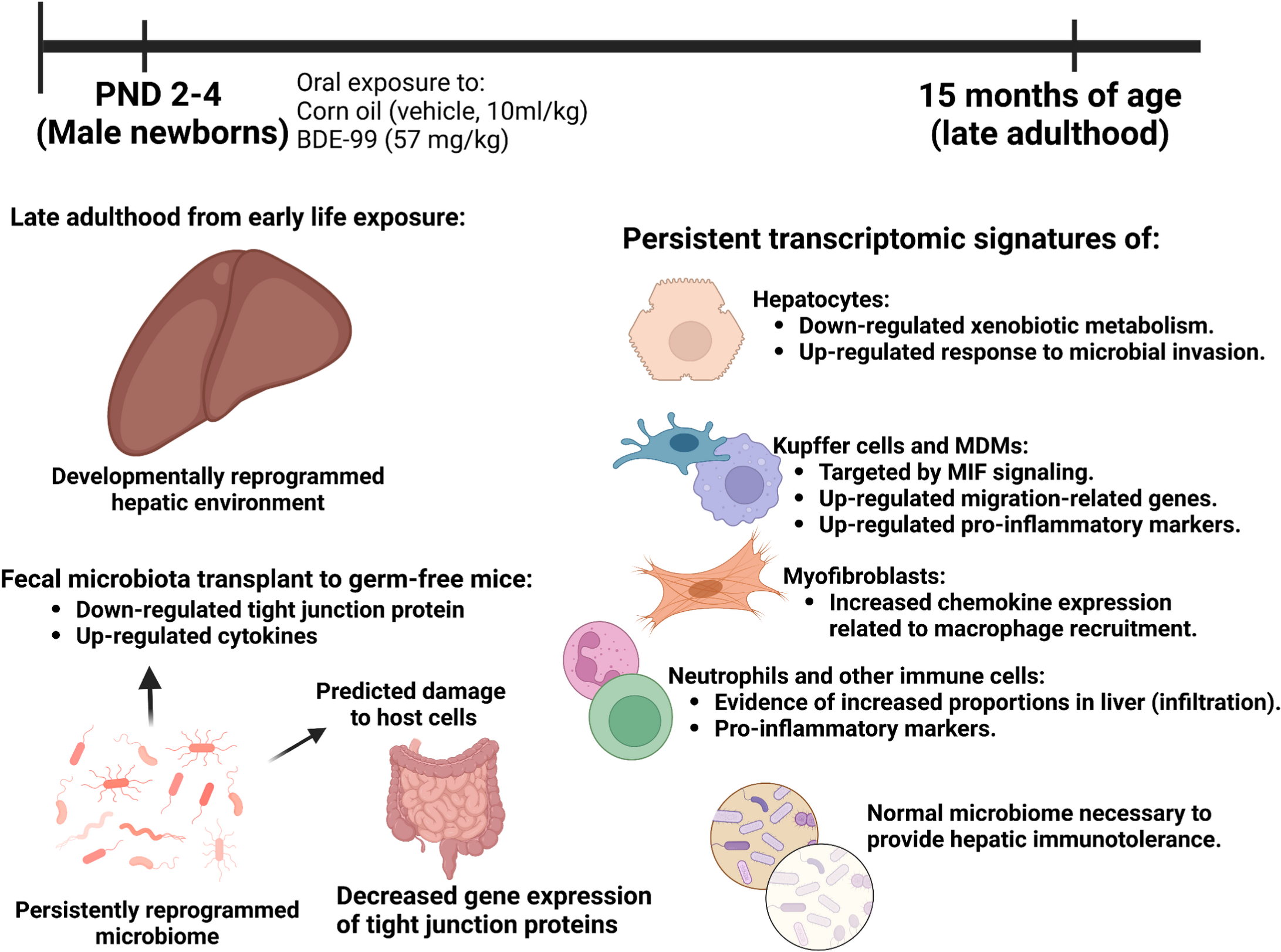
Summary of key findings. Neonatal short-term oral exposure to the human breast milk-enriched persistent organic pollutant, BDE-99, persistently reprogrammed the transcriptome of key liver cell types in late adulthood. Early life exposure to BDE-99 resulted in persistent down-regulation of xenobiotic metabolism and up-regulation of pro-inflammatory signatures in hepatocytes at 15 months of age. The BDE-99 mediated down-regulation of tight junction proteins in the large intestine, together with the dysregulation of liver-disease-associated taxa and the persistently up-regulated hepatic signatures of microbial invasion responses, suggest that gut-liver axis may be a contributor to PBDE-induced alterations in the liver transcriptome. Following neonatal exposure to BDE-99, MIF signaling was persistently increased targeting Kupffer cells and MDMs, which also displayed signatures of inflammatory phenotypes. Chemokines related to macrophage recruitment were persistently up-regulated in myofibroblasts. Early life exposure BDE-99 resulted in increased proportions of neutrophils. The absence of the gut microbiome resulted in increased inflammatory signatures, suggesting that the presence of a healthy microbiome is necessary for the liver to be immunotolerant. In the large intestine, the persistent change in microbial composition in the large intestine is linked to the down-regulation of tight junction proteins and up-regulation of pro-inflammatory cytokines as evidenced by fecal microbiota transplantation. These results suggest that early life exposure to BDE-99 leads to a persistently dysregulated crosstalk between the gut environment and liver and that the persistent effect of neonatal exposure can induce a long-term increased risk for chronic liver disease such as inflammation.

We previously showed a marked sex difference in BDE-99 mediated effects on transcriptome, metabolome, and metagenome within the gut-liver axis, with male adults being more susceptible to the early life BDE-99 exposure-induced changes in hepatic pro-inflammatory and cancer-prone transcriptomic signatures than females (Lim *et al*., 2021). Sex differences are present in response to toxicants, pharmaceuticals, diseases, and developmental reprogramming have been reported (Kim *et al*., 2018; Vyas *et al*., 2019; Deane *et al*., 2001; Kundakovic *et al*., 2013; Lim *et al*., 2021). From our previous study investigating the hepatic signatures following neonatal exposure to BDE-99 in PND 60 adults, we speculated that the differences in the endocrine system and the resulting dissimilarities in hormonal homeostatic state contributed to the sex differences, as the mRNA of androgen receptor was persistently down-regulated only in the young adult males neonatally exposed to BDE-99 (Lim *et al*., 2021). As males were more susceptible to PBDE-mediated hepatic developmental reprogramming than females, investigating whether the higher susceptibility in males continues throughout life into late adulthood is an important toxicological question. Therefore, the present study focused on the male offspring in the investigation of the mechanisms underlying PBDE-developmental reprogramming of the gut-liver axis.

Similar to our earlier report of the effect of neonatal BDE-99 exposure on hepatic reprogramming in young adulthood (PND 60) using bulk RNA-Seq (Lim *et al*., 2021), in the present study we showed that neonatal exposure to BDE-99 also down-regulated many drug-processing genes in hepatocytes in late adulthood (Fig. 3). Because hepatocytes are the main contributor of xenobiotic biotransformation in liver, it is likely that the decreased drug-processing gene expression at PND 60 young adulthood is also a hepatocyte-specific effect. At 15 months of age, the expression of drug-processing genes was increased in non-parenchymal cells by early life BDE-99 exposure; although resident non-parenchymal cells have been reported to carry out xenobiotic biotransformation (Oesch *et al*., 1986; Lafranconi *et al*., 1986), due to the much lower absolute read counts of these genes in these cell populations than in hepatocyte, the overall net effect is expected to be a decrease in drug processing capacity in liver. Together evidence from both PND 60 and 15-month-old male adults showed that BDE-99 mediated decreases in the hepatic expression of drug-processing genes are clearly persistent and may lead to reduced capacity of chemical detoxification over the course of a lifetime. Therefore, neonatal age is a highly sensitive time window for PBDE exposure-mediated prolonged adverse health effects.

Several studies in the literature showed that early life exposure impacts intermediary metabolism in later developmental ages. For example, plasma insulin, glucose sensitivity, and hepatic glutamate dehydrogenase activity were reduced in F1 pups from exposure to DE-71 (an industrial PBDE mixture with BDE-99 ranked among the top most enriched congeners) in mouse dams (Kozlova *et al*., 2020). Increased incidence of liver fibrosis, serum transaminase levels, and lipid peroxidation were reported in mice co-exposed to BDE-47 (another diet and breast milk enriched PBDE congener)and high-fat diet (Yang *et al*., 2019). Furthermore, early-life exposure to BDE-47 persistently up-regulated the lipid influx transporter transcript *Cd36* in 10-month adults in mice (Khalil *et al*., 2018). Our data added to growing evidence of literature showing that early life BDE-99 exposure also impacted xenobiotic biotransformation and inflammation pathways in liver over a time course (present study and (Lim *et al*., 2021). In hepatocytes, we showed that *Cyp1a2*, *Cyp3a11*, and *Cyp4a14*, which are the prototypical target genes for AhR (Vogel *et al*., 2020), PXR (Cui *et al*., 2010), and PPARα (Patsouris *et al*., 2006) were persistently down-regulated by neonatal exposure to BDE-99 in 15-months of age. Because many of these xenobiotic-sensing transcription factors are also involved in intermediary metabolism as implicated in obesity and diabetes (Wang *et al*., 2011; Kerley-Hamilton *et al*., 2012; He *et al*., 2013; Hukkanen *et al*., 2014), the results from our study suggest that the decrease in the signaling of these receptors may also increase the risks of metabolic syndrome in late adulthood. Because aging is another significant risk factor for developing obesity and diabetes, our study linking neonatal PBDE exposure to toxicity in late adulthood is important and calls for attention to factoring early life exposure window into risk assessment in diseases of the elderly.

Exposure to PBDEs is known to promote inflammation in other organs and cell types. For example, chronic exposure to a mixture of three common PBDE congeners (BDE-47, -99, and 209) in human lung cell lines with up-regulated histone modification related to inflammatory genes (Anzalone *et al*., 2021). BDE-47 increased interleukin 7 (IL-7) mRNA in primary normal human bronchial epithelial cells (Anzalone *et al*., 2021) and exacerbated the lipopolysaccharide (LPS)-induced pro-inflammatory response THP-1 macrophages (Longo *et al*., 2021). Low-dose PBDE exposure has also been shown to disrupt genomic integrity and innate immunity in mammary tissue of mice (Lamkin *et al*., 2022). PBDEs increased the expression and secretion of the pro-inflammatory cytokine IL-6 in an immortalized human granulosa cell line (Lefevre *et al*., 2016). BDE-99 reduced the production of IL-1b but increased the production of IL-10 in placental explant cultures (Arita *et al*., 2018). BDE-47-mediated increase in reactive oxygen species induced inflammatory cytokine release from human extravillous trophoblasts *in vitro* (Park *et al*., 2014). In humans, serum BDE-153 was positively associated with alkaline phosphatase and neutrophil counts, which are markers of oxidative stress and inflammation, respectively (Yuan *et al*., 2017). Human peripheral blood mononuclear cells acutely exposed to the commercial PBDE mixture DE-71 showed significantly up-regulated pro-inflammatory cytokines and chemokines (Mynster Kronborg *et al*., 2016). Chronic exposure to BDE-47 resulted in the up-regulation of the nuclear factor erythroid 2 like 2 (Nrf2), which is a key transcription factor of oxidative stress and electrophiles, as well as nuclear factor kappa B (NF-κB) in the mammary tissue in mice (Lamkin *et al*., 2022). F0 rats gavaged with DE-71 throughout pregnancy resulted in increased liver weight and hepatocellular hypertrophy, and F1 rats exposed to DE-71 from postnatal days (PND) 22-42 showed increased body weight and increased splenic T cell populations (Bondy *et al*., 2013). These results in the literature have suggested that inflammation is a key mechanism for PBDE-mediated toxicity.

One key finding from our study is the persistently up-regulated hepatic immune response signatures at 15 months of age following neonatal exposure to BDE-99 (Fig. 4). Our results align with previous findings of early life exposure and dysregulated immune responses. The inflammatory marker, signal transducer, and activator of transcription 3 (*Stat3*) was up-regulated in F1 male offspring by DE-71 exposure to F0 dams (Kozlova *et al*., 2022). Early life exposure to polychlorinated biphenyl (PCB) 126, another persistent organic pollutant, when combined with high-fat diet, persistently up-regulated serum inflammatory cytokines (Tian *et al*., 2022). We showed that there was a tendency towards a persistent increase in hepatic macrophage populations (i.e. Kupffer cells and MDMs) and dendritic cells as well as a significant increase of neutrophil translocation to the liver by neonatal exposure to BDE-99 (Fig. 4A). Fatty acids related to anti-inflammatory processes (Jia *et al*., 2020; Saresella *et al*., 2020; Rial *et al*., 2016) were persistently down-regulated in both serum and liver (Fig. 4B-4C). Single cell transcriptomic data in liver suggested a persistent increase in response to LPS and other microbial constituents in hepatocytes of adults neonatally exposed to BDE-99 (Fig. 4D). The influx of certain microbial products, such as LPS, leads to inflammatory liver injury (Zamani et al. 2013; Longo et al. 2021; Colman 2015). Lipopolysaccharide (LPS) can bind to binding proteins in plasma, which is delivered and sensed by CD14 (Zamani *et al*., 2013). Following neonatal exposure to BDE-99, the expression of *Cd14* was persistently up-regulated in MDMs, further providing evidence of increased influx of microbial products. Chemokines and cytokines that act as chemoattractors of immune cells and pro-inflammatory mediators in the liver were persistently up-regulated in various cell types following neonatal exposure to BDE-99 (Fig. 4E). Specifically, in myofibroblasts, the pro-inflammatory cytokine Il18, as well as macrophage chemoattractors Ccl2 and Ccl7 were persistently up-regulated by neonatal exposure to BDE-99 (Gschwandtner, Derler, and Midwood 2019; Xie et al. 2021). Overall, our results suggest that early life exposure to BDE-99 promotes the liver to be in a pro-inflammatory state.

Inflammatory responses are the products of delicate communications across cells (Altan-Bonnet and Mukherjee 2019; Ramesh, MacLean, and Philipp 2013; Proudfoot 2002). These altered signaling and inflammatory attacks remain as key mechanisms for cell damage and alterations in hepatic functions. For instance, hepatic macrophage populations, upon infection, can secrete IL6 and IL-1β, which can induce the proliferation of cholangiocytes and reactions in ductal cells (J. Park et al. 1999; Xiao et al. 2015). Through growth factors, such as transforming growth factor-beta 1 (TGF-β), hepatic macrophages can activate hepatic stellate cells, leading to myofibroblasts and fibrogenesis in the liver (Vannella and Wynn 2017). These results point out the importance of cellular signaling and the roles of macrophages in hepatic disease progression. We showed that hepatic macrophage populations were developmentally reprogrammed (Fig. 5). From early life exposure to BDE-99, hepatic macrophages, overall, showed pro-inflammatory signatures and up-regulated markers, such as Cxcl10 and Il6 (Fig. 5C). CXCL10 functions as a chemoattractant of inflammatory cells and regulates cell growth (Liu, Guo, and Stiles 2011). In the liver, CXCL10 activates CXCR3 expressing leukocytes, such as T cells and NK cells, thereby promoting pro-inflammatory cascades in the liver, and is closely related to IL6-mediated inflammation (Liu, Guo, and Stiles 2011; Hintermann et al. 2010; Malik and Ramadori 2022). Phenotypically, we observed a mild increase in hepatic inflammation likely due to circulating inflammatory cells (Fig. S8). Interestingly, the macrophages were targeted by predicted up-regulation in MIF signaling (Fig. 5). One key mediator in the up-regulation of MIF signaling was from hepatocytes (Fig. 5). Exposure to ethanol in hepatocytes up-regulated and released MIF, and chimeric mice that expresses MIF in non-myeloid cells, including hepatocytes, up-regulated pro-inflammatory cytokines and chemokines (Marin et al. 2017). Non-myeloid cell-specific MIF knockout mice were protected from ethanol-induced liver injury, including inflammation signatures (Marin et al. 2017). These results suggest that hepatocyte-derived MIF signaling may be an important mechanism for the persistent up-regulated immune signatures following early life exposure to BDE-99.

Leaky gut is closely linked to and often accompanied by dysbiosis due to mechanisms including imbalanced nutrient absorption, altered mucus layer, and impairment of microbial sensing in host cells (Lobionda *et al*., 2019; Kinashi and Hase, 2021; Stolfi *et al*., 2022). Numerous studies have shown the link between dysbiosis and liver disease progression, such as in alcoholic liver disease (Hartmann *et al*., 2015; Morencos *et al*., 1995; Hartmann *et al*., 2013), non-alcoholic fatty liver disease (Panasevich *et al*., 2017; Raman *et al*., 2013), non-alcoholic steatohepatitis (Brandl and Schnabl, 2017; Carter *et al*., 2021), and liver cancer (Zhang *et al*., 2019; Schneider *et al*., 2022; Bartolini *et al*., 2021). The persistently up-regulated MIF signaling in hepatocytes and the resulting pro-inflammatory signatures in macrophages, as well as other cell types including neutrophils, may be from the persistently dysregulated gut environment following early life exposure to BDE-99. Furthermore, significantly up-regulated neutrophils and a trend towards increased T and NK cell populations in the liver suggest persistently up-regulated influx of inflammatory immune cells (Fig. 4). Together the trend of increased myofibroblast populations and up-regulated macrophage attractants indicate a possible mechanism for the persistently up-regulated pro-inflammatory signatures by early life exposure to BDE-99. In addition to the up-regulation of genes involved in the influx of microbial products in hepatocytes, the persistently decreased levels of fatty acids involved in anti-inflammation in serum and liver are also byproducts of the gut microbiota (Rios-Covian *et al*., 2020; Gregor *et al*., 2021; Jia *et al*., 2020). This is likely due to a persistent increase in gut permeability, as evidenced by a persistent decrease in tight junction protein transcripts in the large intestine in late adulthood. In the present study, we show that early life exposure to BDE-99 persistently down-regulated transcripts of tight junction proteins in the large intestine (Fig. 6A). The intestinal barrier functions as an important regulator for the outflux of microbial products and fragments, and the up-regulated intestinal permeability is closely linked to inflammation (Farré *et al*., 2020; Fukui, 2016).

In late adulthood, neonatal exposure to BDE-99 lead to a persistently dysbiotic gut microbiome. From the initial BDE-99 exposure that occurred early in life, 43 taxa at the full species level were persistently dysregulated. Multiple beneficial and commensal taxa were persistently down-regulated in the large intestine by early life exposure to BDE-99. For example, *Blautia obeum* has been investigated as a potential probiotic (Liu *et al*., 2021). *Faecalibacculum rodentium* is an endogenous murine microbiota member associated with anti-inflammation (Zagato *et al*., 2020). *Mediterraneibacter faecis* is dominant in healthy human gut than moderate malnutrition state (Kamil *et al*., 2021). *Akkermansia muciniphila* is generally considered as beneficial and anti-inflammatory (Raftar *et al*., 2022; Zheng *et al*., 2022; Zhang *et al*., 2021), *Dorea longicatena* is common gut commensal microbiota (Schirmer *et al*., 2016). Furthermore, taxa that are negatively associated with diseases were also persistently down-regulated by neonatal exposure to BDE-99. For instance, *Harryflintia acetispora* is inversely correlated in rheumatoid arthritis-mediated inflammation in humans (Lee *et al*., 2019). *Parvibacter caecicola* is associated with anti-tumor processes (Li *et al*., 2020; Routy *et al*., 2018). *Dubosiella newyorkensis* is beneficial taxon with anti-aging properties in mice (Liu *et al*., 2023). *Anaerotruncus colihominis* is negatively correlated with experimental multiple sclerosis in mice (Bianchimano *et al*., 2022). The persistently up-regulated taxa by early life exposure to BDE-99 were associated with diseased states and disease development. *Erysipelatoclostridium ramosum* was increased in Crohn’s disease in humans (Sankarasubramanian *et al*., 2020) and was decreased in Crohn’s treatment Metronidazole + Azithromycin in humans (Mah *et al*., 2023). Campylobacter hyointestinalis is an infectious microbe that displays acute inflammation (Igwaran and Okoh, 2019; Facciolà *et al*., 2017). *Anaerotignum lactatifermentans* was increased in pelvic irradiation-induced intestinal mucositis in mice (Segers *et al*., 2021), as well as in typhlitis in turkey (Abdelhamid *et al*., 2021), *Lactobacillus intestinalis* was increased in spinal cord injury in rats (O’Connor *et al*., 2018), as well as in type 2 diabetes in rats and humans (Wang *et al*., 2020; Sato *et al*., 2014). *Agathobacter rectalis* was increased in ulcerative colitis in humans (Lavelle *et al*., 2022) and in insulin resistance syndrome in humans (Álvarez-Mercado *et al*., 2019). *Enterocloster citroniae* was positively correlated with type 2 diabetes in humans (Ruuskanen *et al*., 2022). These results show that early life exposure to BDE-99 altered the gut microbiome that harbors more diseased-associated microbiota while decreasing beneficial and commensal taxa.

In addition to the persistently altered gut microbiota, caproic acid is a medium-chain fatty acid that is involved in pro-inflammatory processes by promoting Th1 and Th17 differentiation (Saresella *et al*., 2020). We found that caproic acid was persistently up-regulated in the large intestinal content and large intestine, which provides evidence for a causal relationship between dysregulated gut microbiota and increased abundance of caproic acid in the gut environment. Recently, our studies and other DOHaD investigations on PBDE-mediated developmental reprogramming reported persistent alteration in the gut environment. For example, we showed that perinatal exposure to BDE-47 resulted in persistent dysbiosis in adult mice (Gomez *et al*., 2021). Perinatal exposure to BPA persistently dysregulated the microbiome in adult rabbits (Malaisé *et al*., 2017; Reddivari *et al*., 2017). We showed that the gut microbiome was persistently dysregulated by neonatal BDE-99 exposure at PND 60 (Lim *et al*., 2021). Our results add another layer of interesting complexity to the role of the altered microbiota from toxic exposures that occur during critical developmental windows, and the altered risk and delayed onset of disease development.

To validate the role of the microbiome in regulating hepatic and gut immune responses, we utilized germ-free mice. In the absence of microbiome (i.e. deviation from a normal gut environment), pro-inflammatory complement signaling was predicted to be up-regulated targeting hepatic macrophages and DCs (Fig. 7B). In addition, the absence of microbiome was accompanied by up-regulated *Cxcl12,* which activates leukocytes and results in pro-inflammatory responses (Dotan *et al*., 2010; García-Cuesta *et al*., 2019). Pro-inflammatory cytokines including *Tnfα*, *Il-1α*, and *Il-5*, as well as regulators of inflammation, such as *Vegf1*, *Icam1*, and *Egr1* (Yan *et al*., 2021; Kong *et al*., 2018; Bui *et al*., 2020) were up-regulated in the absence of microbiome (Fig. 7C). These results lead us to the conclusion that the presence of a normal microbiome is necessary in priming the liver to be immunotolerant. Fecal microbiota transplantation of the large intestinal content to germ-free mice was used to validate the role of persistent dysbiosis in regulating gut health from early life exposure to BDE-99. We showed that the transplantation of large intestinal content from adults neonatally exposed to BDE-99 resulted in a persistent down-regulation of both tight junction proteins in the large intestine (Fig. 8). These results suggest that the altered taxa in the gut was sufficient to disentangle the gaps in the gut epithelium and can increase leakage of microbial products that may impact the homeostatic state in other organs, such as the liver. IL22, which can function as an anti-inflammatory cytokine, is a marker of Th22 cells, and its the main function is to protect epithelial barriers and can act as a modulator of inflammation at sites of injury to promote wound healing (Eyerich and Eyerich, 2015; Keir *et al*., 2020). IL17 can act as a pro-inflammatory cytokine that can be produced by Th17 cells (Tesmer *et al*., 2008; Singh *et al*., 2014) that can act as a chemoattractor of inflammatory immune cells, such as neutrophils (Zenobia and Hajishengallis, 2015). Our results showed that fecal microbiota transplant of the large intestinal content of adults neonatally exposed to BDE-99 tended to decrease the expression of *Il22*, while up-regulating the expression of *Il17* and *Tnfα*, suggesting that the change in microbial composition by early life exposure to BDE-99 leads to a pro-inflammatory gut environment. With the persistently up-regulated functional prediction related to cytotoxic processes by up-regulated gut microbes (Fig. 6C), our results suggest that early life exposure to BDE-99 promotes the gut environment to be in a pro-inflammatory state, which may aggravate the responses observed in the liver. Overall, our findings suggest the gut-liver axis is an important mechanism in regulating cell type-specific developmental reprogramming related to immune response signatures.

The present study has limitations, including a lack of protein and enzyme activity quantifications, and small sample sizes for histology to make an accurate phenotypic assessment for hepatic inflammation. Moreover, most cell type proportions were not statistically different between the vehicle and BDE-99 groups due to the limited sample sizes used for scRNA-seq. To overcome these limitations, protein expression by immunohistochemistry in the livers can be performed to validate the persistently up-regulated pro-inflammatory signatures by neonatal exposure to BDE-99. The dose of BDE-99 may not be a realistic dose for humans. However, previous screening results found that the serum concentrations of BDE-99 in humans are between 1.3 and 290 pg/g (Frederiksen *et al*., 2009). As BDE-99 is enriched in breast milk, newborns are more vulnerable to BDE-99 exposure and are exposed to BDE-99 at approximately 5.7 ng/g liquid weight (Lorber, 2008). A factor of 10 was applied to extrapolate inter-species differences, which lead to the concentration of BDE-99 used in this study. The dose of BDE-99 is also comparable to other toxicological investigations of acute BDE-99 exposure (Li *et al*., 2017, 2018; Scoville *et al*., 2019), as well as our previous study that investigated the persistent dysregulation in the gut-liver axis at PND 60 by neonatal BDE-99 exposure (Lim *et al*., 2021). Although we provide multiple evidence of an increased influx of microbial molecules following early life exposure to BDE-99 and persistent alterations in the gut environment, tight junctions at the protein level were not directly measured. Quantification of protein expression can be done in the intestines, and microbial DNA and microbial molecules, such as LPS, can be quantified in the liver.

Despite its limitations, our present study is the first to show that neonatal exposure to BDE-99 reprograms the liver in a cell type-specific manner in late adulthood and suggests that the gut-liver axis mediated liver inflammation is a possible mechanism for delayed onset of liver injuries. By characterizing the persistent changes in liver cell networks using single cell transcriptomics, our study contributes to precision medicine in vulnerable aging populations who are at higher risk of exposure to environmental contaminants at sensitive time windows.

## ACKNOWLEDGEMENT

This work was supported by NIH grants R01ES019487, and R01ES025708, the University of Washington Interdisciplinary Center for Exposures, Diseases, and Environment (EDGE) (P30ES007033), Environmental Pathology/Toxicology Training Program (T32ES007032), Environmental Health and Microbiome Research Center (EHMBRACE), and the Sheldon Murphy Endowment.

