## Supplemental figure captions for "Single cell transcriptomics unveiled that early life BDE-99 exposure reprogrammed the gut-liver axis to promote a pro-inflammatory metabolic signature in male mice at late adulthood"

**Figure S1** UMAP representation of cell types in adults neonatally exposed to vehicle (left) or BDE-99 (right).

**Figure S2.** Gene ontology enrichment of all expressed drug processing genes in hepatic cell types. Cell type-specific conserved differentially expressed genes present in both the vehicle and BDE-99 cells were used (Bonferroni-adjusted *p*-value < 0.05). Columns represent cell types, rows indicate individual genes (not shown), left clusters show k-means clustering results, and right text show gene ontology enrichment results for each cell type-enriched drug processing genes. Yellow and blue show high and low expression, respectively.

**Figure S3.** Top 10 up-regulated (**A**) and down-regulated (**B**) gene ontology enrichment results in endothelial cells, cholangiocytes, stellate cells, and myofibroblasts in adult mouse livers following neonatal exposure to BDE-99. Dotted lines represent $-\log_{10}$ FDR-adjusted *p*-value at 0.05.

**Figure S4.** Log2 fold change values in endothelial cells, cholangiocytes, stellate cells, and myofibroblasts for persistently dysregulated drug processing genes relative to hepatocytes in the vehicle (**A**) and BDE-99-exposed group (**B**). Red and blue in the heatmap represent the $\log_2$ fold change of liver genes of the BDE-99 exposed adults in non-parenchymal cells with respect to hepatocytes. "Direction" indicates whether a gene is up- or down-regulated from neonatal exposure to BDE-99 (Bonferroni-adjusted *p*-value < 0.05).

**Figure S5.** Top 10 up-regulated (**A**) and down-regulated (**B**) gene ontology enrichment results in cDC and pDC in adult mouse livers following neonatal exposure to BDE-99.

**Figure S6.** Top 10 down-regulated gene ontology enrichment results in Kupffer cells and MDMs in adult mouse livers following neonatal exposure to BDE-99. Dotted lines represent $-\log_{10}$ FDR-adjusted *p*-value at 0.05.

**Figure S7.** Top 10 up-regulated (**A**) and down-regulated (**B**) gene ontology enrichment results in myofibroblasts in adult mouse livers following neonatal exposure to BDE-99. Dotted lines represent -$\log_{10}$ FDR-adjusted *p*-value at 0.05.

**Figure S8. A.** H&E staining of results for males neonatally exposed to vehicle or BDE-99. The left and right holes represent the central vein and portal triad, respectively. Kupffer cells and lymphocytes are located in the sinusoids. **B.** Pathology evaluation results showing the incidence of bile duct hyperplasia (yellow arrows) and immune infiltration (red arrows) by neonatal exposure to BDE-99.

**Figure S9.** UMAP representation of cell types in conventional (left) and germ-free (right) mouse livers.

**Figure S10.** Top 10 up-regulated (higher in germ-free cell types) gene ontology terms in hepatocytes, cholangiocytes, and myofibroblasts (**A**) and in Kupffer cells and MDMs (**B**). Dotted lines represent -$\log_{10}$ FDR-adjusted *p*-value at 0.05.

**Table S1.** Cell type-specific marker genes.

**Table S2.** Cell type-specific persistently differentially expressed genes by neonatal exposure to BDE-99.

**Table S3.** Direction of regulation for persistently dysregulated drug processing genes by neonatal exposure to BDE-99.

**Table S4.** log2 fold change of persistently dysregulated drug processing genes in vehicle group relative to the expression of hepatocytes.

**Table S5.** log2 fold change of persistently dysregulated drug processing genes in BDE-99 group relative to the expression of hepatocytes.

**Table S6.** Cell type-specific differentially expressed genes comparing conventional and germ-free male adult mouse liver.

**Table S7.** List of primer sequences used for RT-qPCR.
