## Supplementary figures and images for "Single cell transcriptomics unveiled that early life BDE-99 exposure reprogrammed the gut-liver axis to promote a pro-inflammatory metabolic signature in male mice at late adulthood"

### Supplemental figures

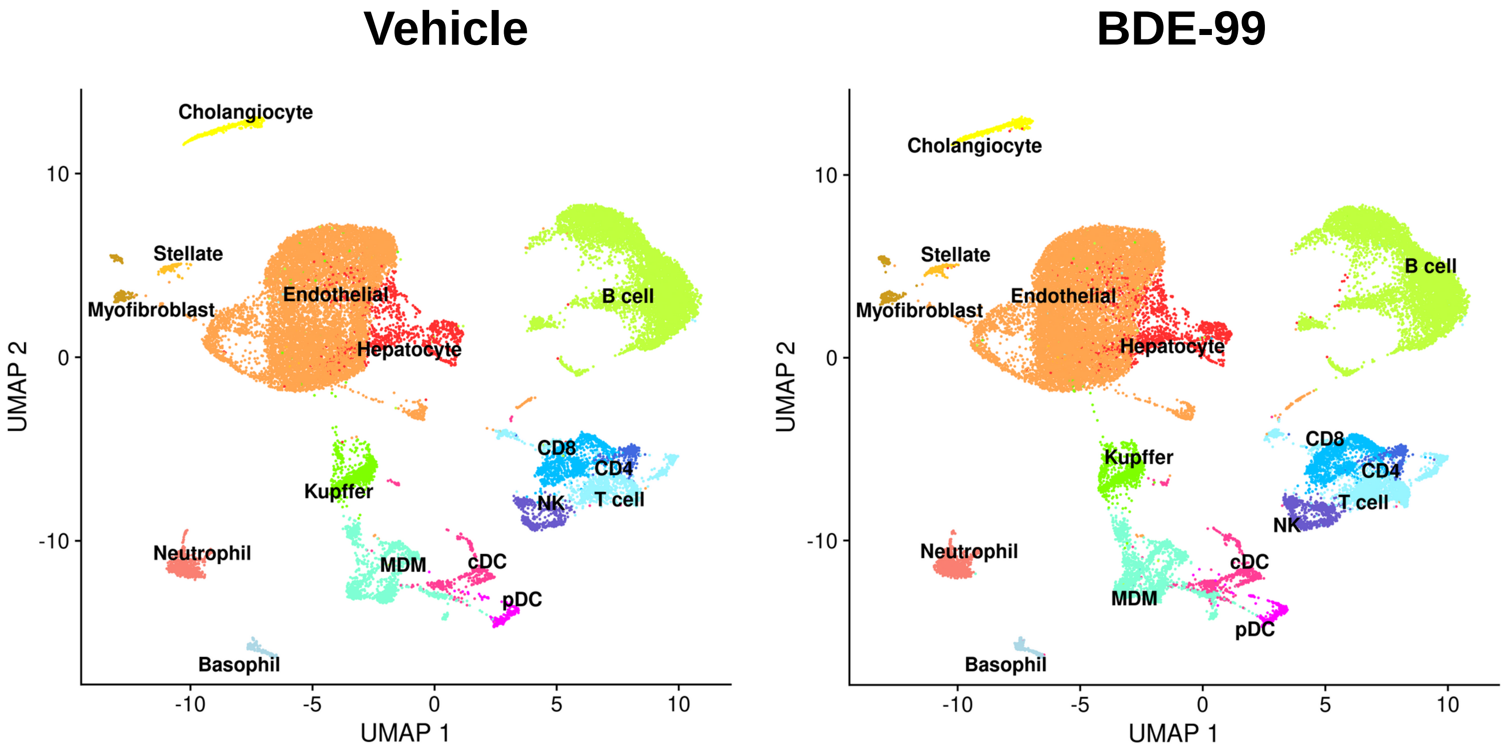

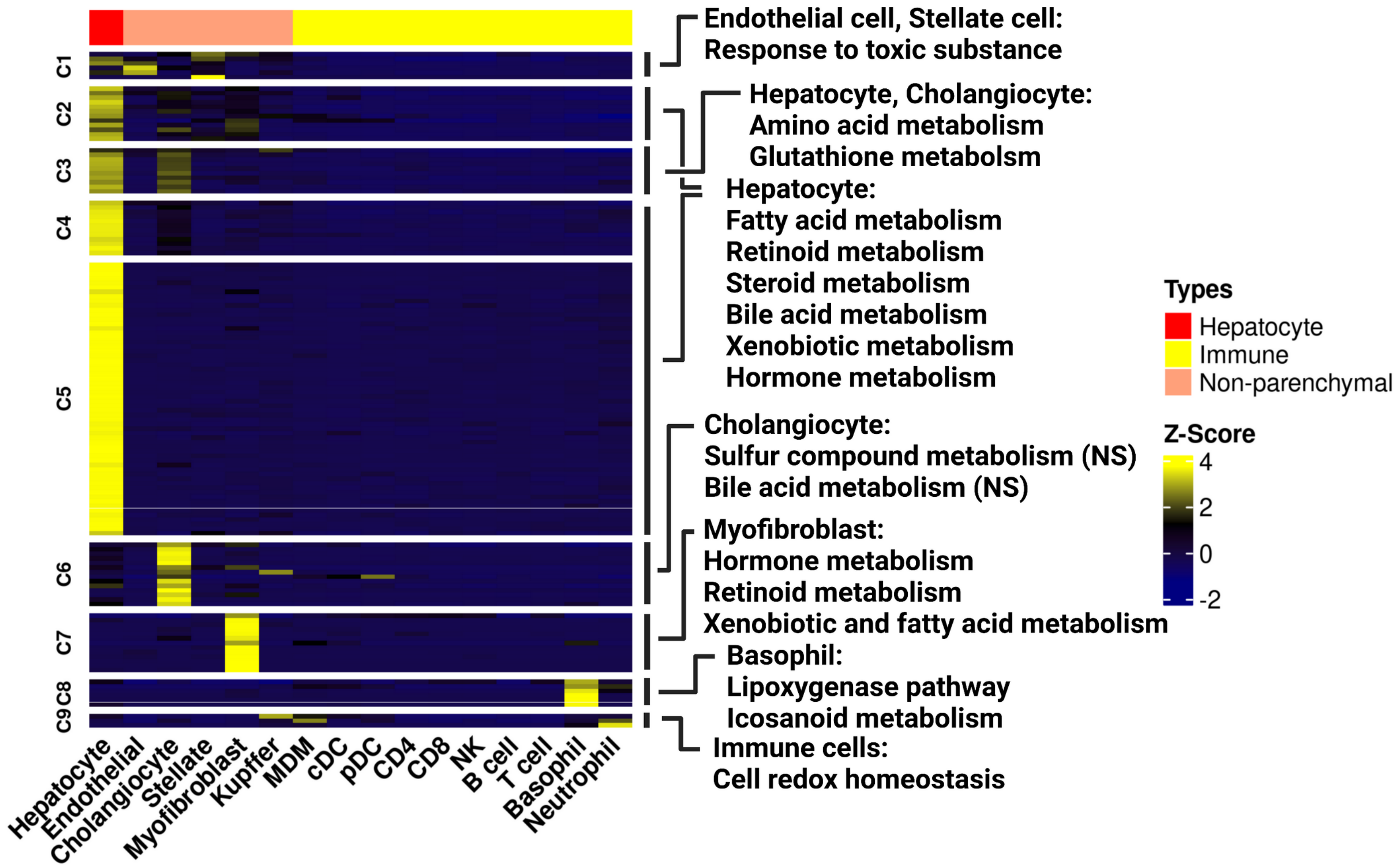

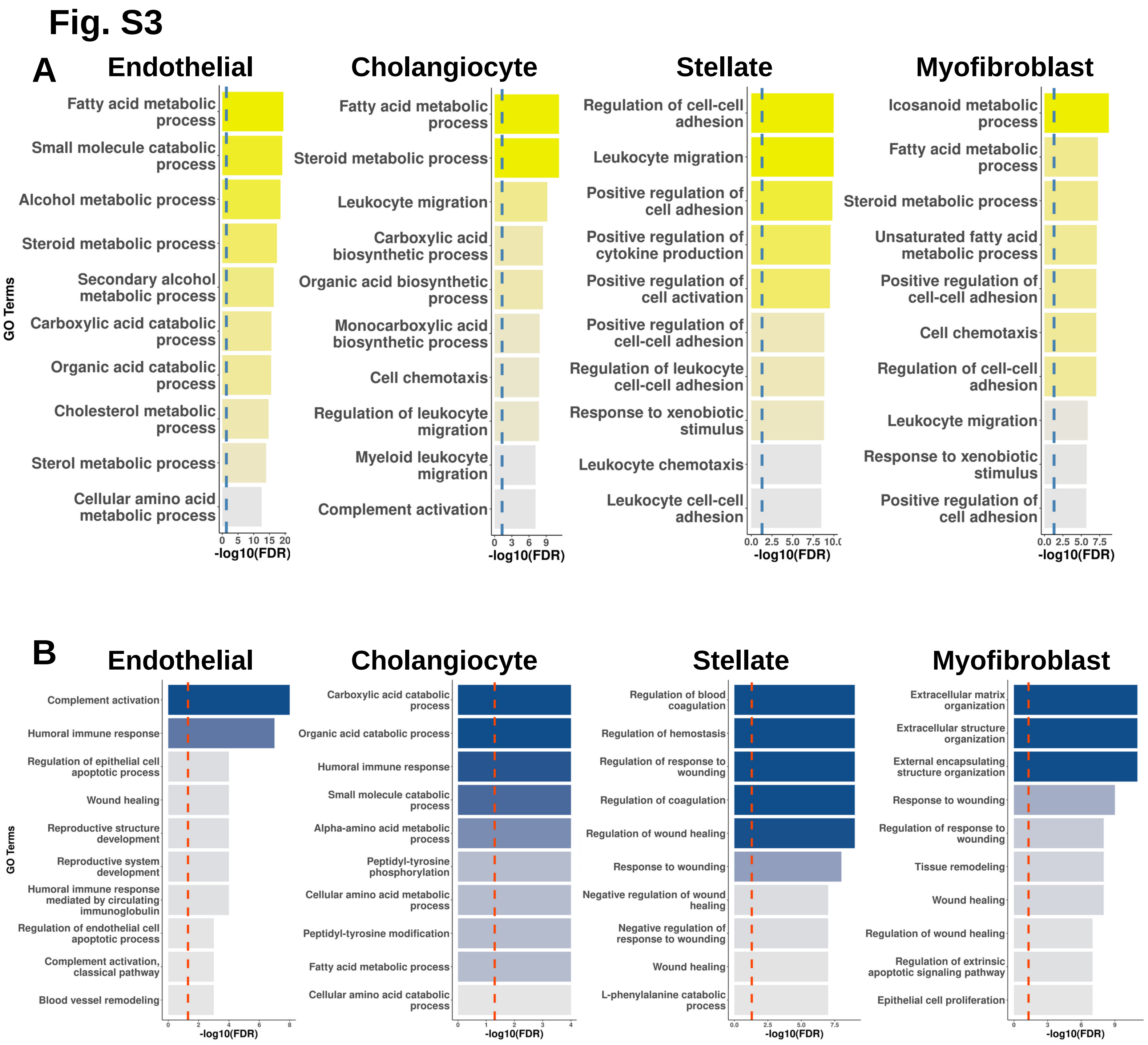

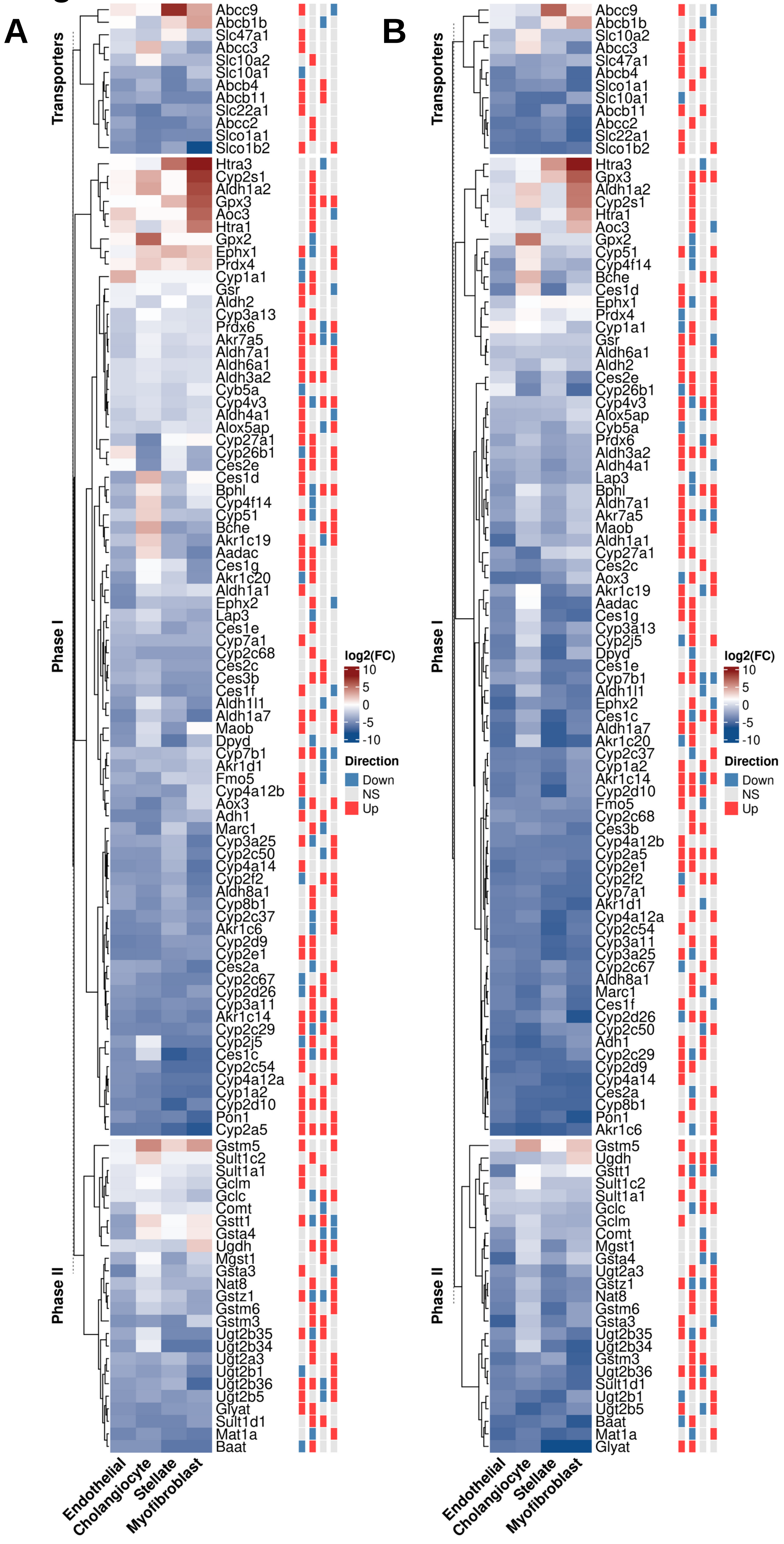

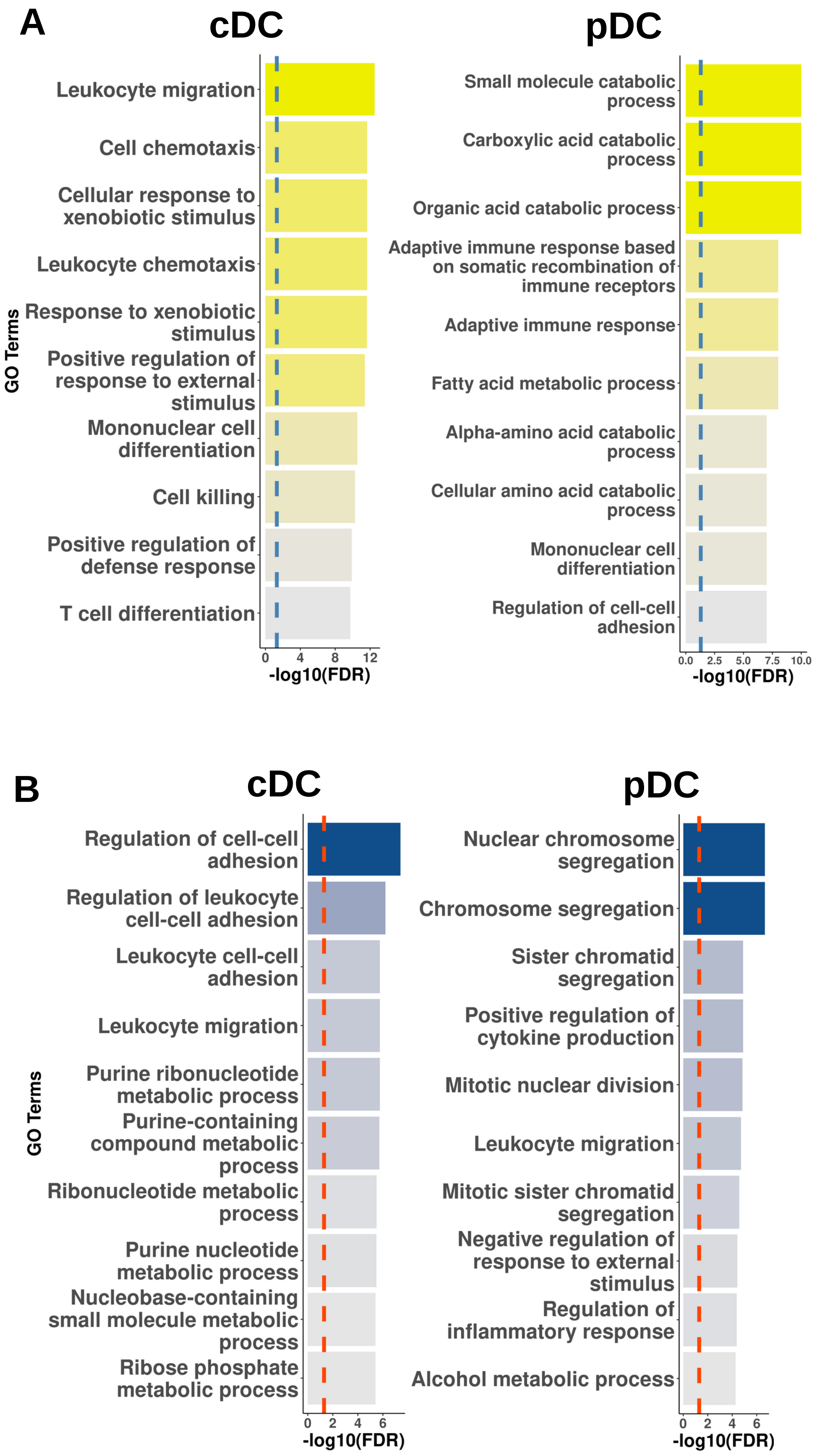

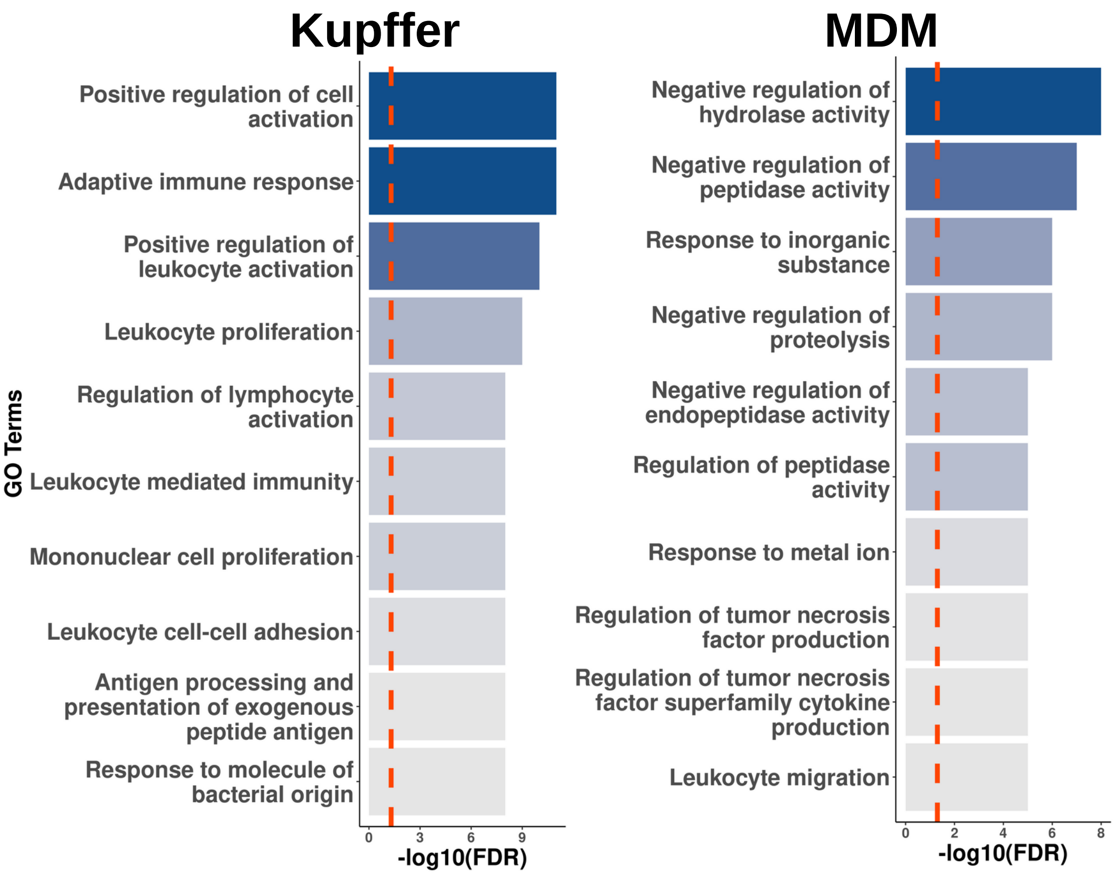

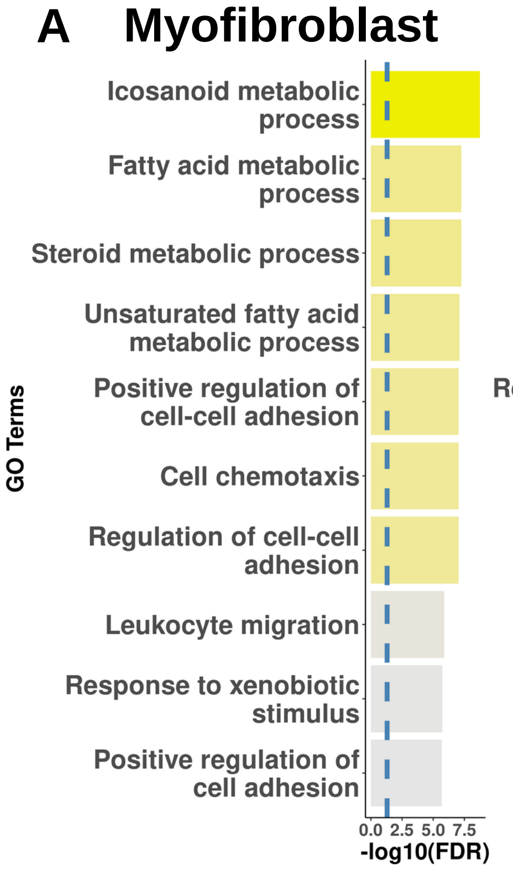

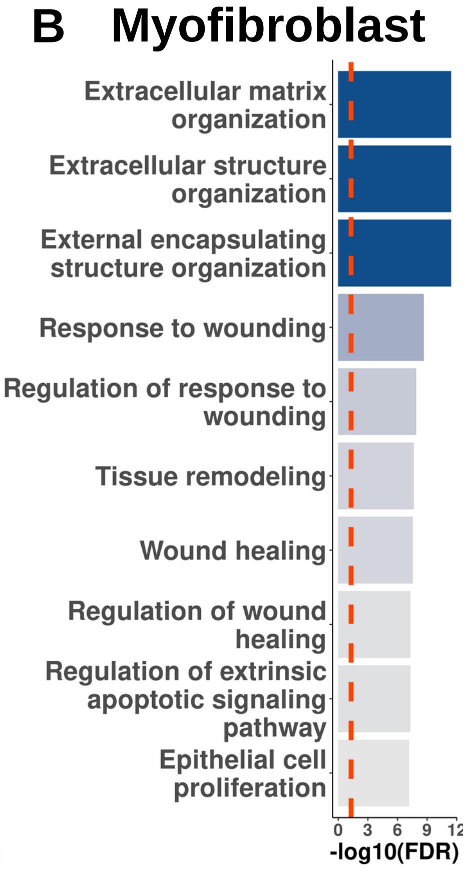

#### Fig. S8

**A**

**Vehicle** **BDE-99**

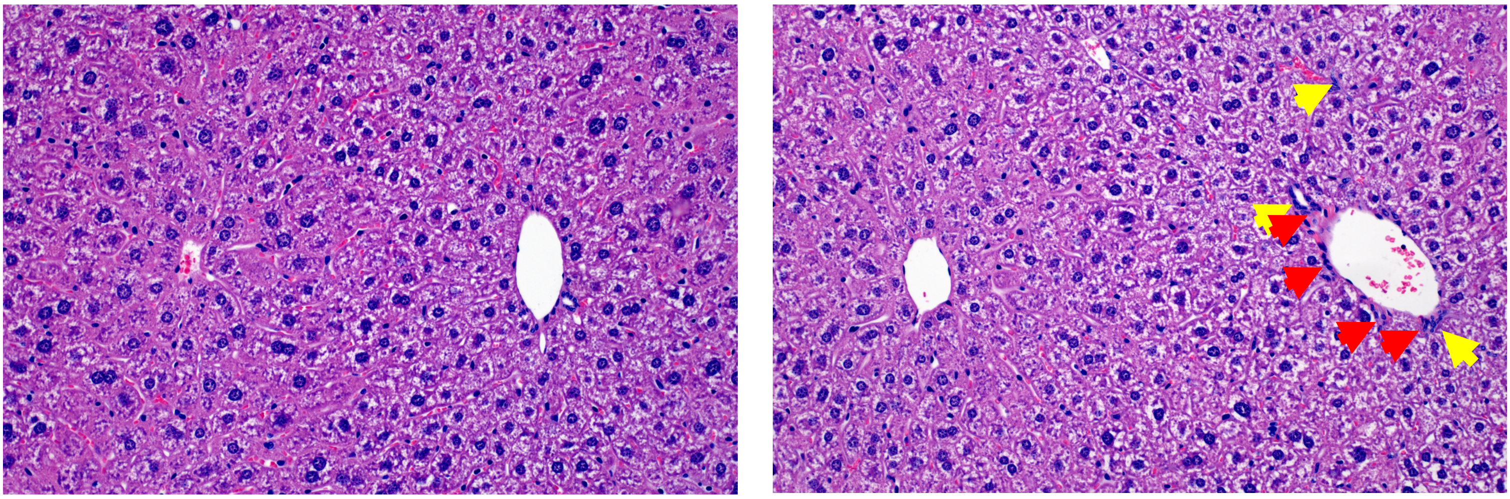

**B**

| | Incidence | |
|---|---|---|
| | **Bile duct hyperplasia** | **Immune infiltration** |
| **Vehicle** | 0/3 | 0/3 |
| **BDE-99** | 1/5 | 1/5 |

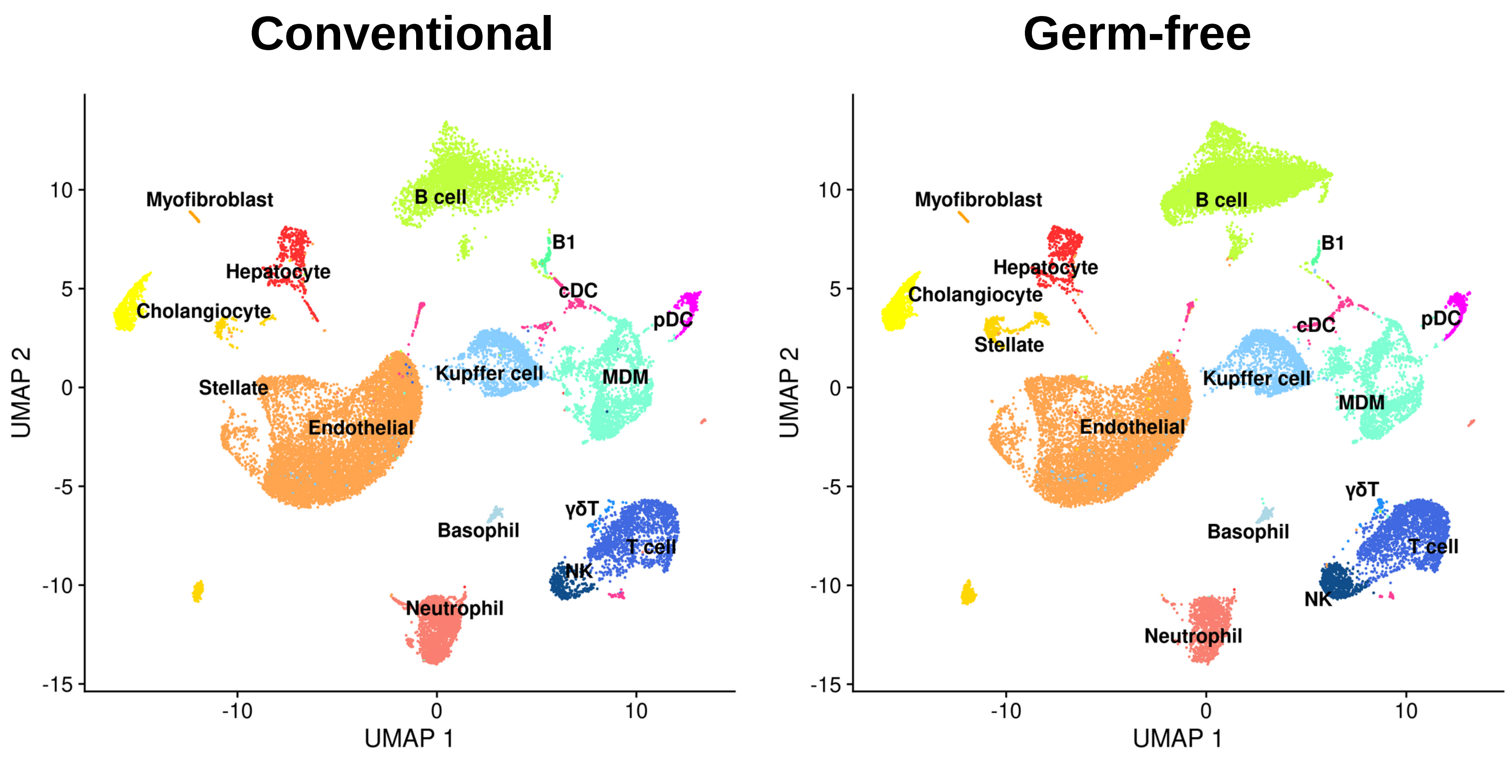

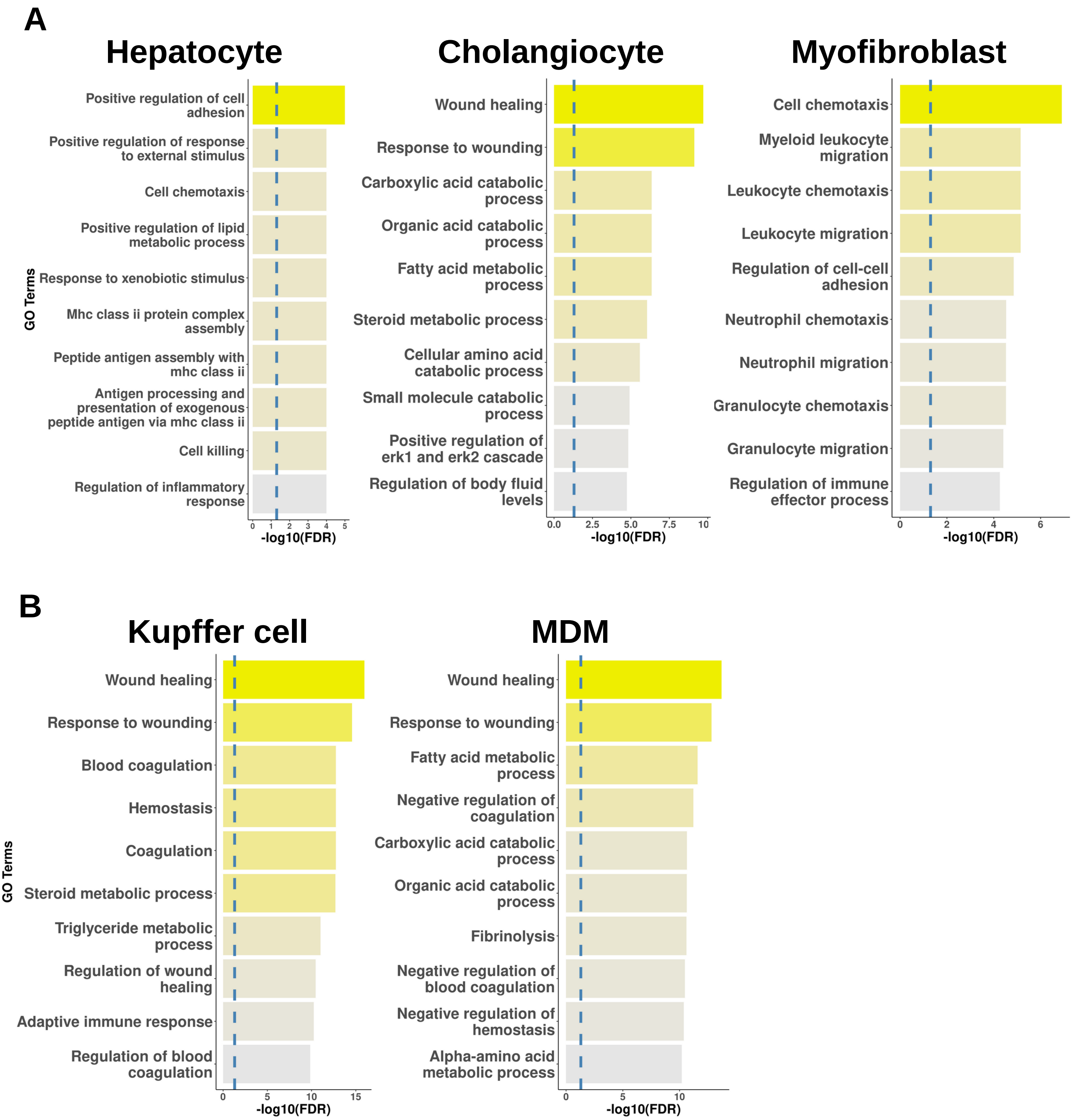
